# Hierarchical and Context-Dependent Encoding of Actions in Human Posterior Parietal and Motor Cortex

**DOI:** 10.1101/2025.11.10.687245

**Authors:** V. Bougou, J. Gamez, E. R. Rosario, C. Liu, K. Pejsa, A. Bari, R. A. Andersen

**Affiliations:** Division of Biology and Biological Engineering, California Institute of Technology, Pasadena, CA; T&C Chen Brain-machine Interface Center, California Institute of Technology, Pasadena, CA; Casa Colina Hospital and Center for Healthcare, Pomona, CA; Department of Neurological Surgery, Keck School of Medicine of USC, Neurological Surgery, Los Angeles, CA; Department of Neurological Surgery, University of California, Los Angeles, CA

## Abstract

Action understanding requires internal models that link vision to motor goals. In monkeys, mirror neurons demonstrate motor resonance during observation, but single-unit evidence in humans is limited, leaving open whether such representations rely solely on motor resonance. We recorded neural activity from motor cortex (MC) and superior parietal lobule (SPL) in two tetraplegic participants implanted with Utah arrays while they intended or observed hand actions. MC strongly encoded intention but showed only weak, feature-specific overlap during observation, evident primarily at the population level. SPL, in contrast, supported shared models across intended movement and observation formats at both single-unit and population levels. In variants with incongruent instructed and observed actions, SPL encoded observed actions only when behaviorally relevant, whereas MC remained intention-dominant. Our results identify a context-dependent gating mechanism in SPL and suggest a hierarchical organization in which MC maintains intention-specific codes while SPL flexibly links observed input with internal goals to support action understanding.

## Introduction

The ability to interpret the actions of others, to identify what is being done, by whom, and in what context, is a fundamental feature of intelligent behavior. It relies on internal representations that can flexibly map visual input onto motor knowledge, enabling the brain to simulate, predict, and interpret observed movements. For decades, these processes have been framed through the lens of mirror mechanisms: the idea that shared neural codes support both action execution and observation^1–3^. Yet the scope, structure, and cortical distribution of these representations remain deeply debated^4^. Is action understanding underpinned by motor-constrained reactivations of one’s own movements, or by more abstract, generalizable codes that transcend the observer’s motor repertoire? And how are these representations organized across regions traditionally implicated in sensory-motor control versus those thought to support higher-order cognitive functions?

Mirror neurons, first identified in macaques, respond during both execution and observation, supporting the motor simulation theory that action perception relies on covert motor reactivation ^1,2,5–8^. While influential, this framework has been refined by evidence that mirror responses are shaped by prior experience and task context (contextual flexibility), that individual units often encode only subsets of action features (partial tuning), and that visuomotor congruence emerges more robustly at the ensemble rather than single-cell level (population encoding). ^9^. These findings point to more complex and heterogeneous mechanisms than strict motor mirroring can explain. Converging theoretical work suggests that cognitive flexibility emerges from internal models that are compositional, building complex structures from separable and recombinable elements, and generalizable capturing abstract structure that extends across contexts and tasks to support adaptive behavior.^10–12^ Yet in humans, direct single-neuron evidence remains scarce, and the broader principles underlying action representations, particularly their flexibility and generalizability, are still not well defined.

These theoretical shifts, from rigid mirroring to flexible, predictive, and context-sensitive encoding, highlight the need to reassess how action representations are structured in the human brain. Action observation provides a unique lens into this flexibility. Unlike execution, it allows testing whether neural populations encode actions disentangled from the specific motor commands or output normally coupled to them This is particularly relevant in higher-order regions such as the parietal cortex, where mixed selectivity is prevalent^13–16^ and encoding may reflect abstract and goal-directed representations rather than motor-specific plans^17–19^. Assessing such generalizability requires moving beyond single-neuron selectivity towards the structure of population codes. Recent advances in neural manifold analysis^20^ and representational geometry ^21^ (the structure of neural population activity, often captured as trajectories) have shown that population activity can reveal compositional and generalizable subspaces, but these approaches have rarely been applied to investigate action representations^22^. To date, only a handful of studies have examined action observation responses at the single-neuron level in humans^22–24^, and none have systematically characterized the geometry or cross-format alignment, i.e. generalizatrion, of action representations in core regions of the motor system.

To investigate how intended and observed actions are represented in the human brain, we recorded single– and multi-unit activity together with local field potentials from motor and posterior parietal cortices in two tetraplegic participants implanted with Utah arrays. Using a factorial design that manipulated hand, action type, and movement direction, we compared neural representations across intention and observation. Our analyses revealed a gradient of representational overlap, with posterior parietal cortex encoding action identity in a format-general manner across intention and observation, whereas motor cortex representations were predominantly intention-specific, with observation responses reflecting only a latent projection of the intention-related structure. To test whether this overlap was modulated by behavioral relevance, we introduced a dissociation task that decoupled instructed and observed actions. This manipulation demonstrated that parietal representations of observed actions emerged only when those actions were behaviorally relevant, consistent with a gating mechanism driven by task demands, whereas motor cortex consistently reflected only the instructed movement. Together, these findings provide direct human electrophysiological evidence that action representations extend beyond motor mirroring, supporting a hierarchical and context-dependent coding framework in the frontoparietal system.

## Results

We recorded single– and multi-unit activity as well as local field potentials (LFPs) from two tetraplegic participants (JJ, RD) implanted with Utah arrays targeting motor and posterior parietal cortices. Participant JJ had 96-channel arrays in the hand knob area of the primary motor cortex (MC) and the superior parietal lobule (SPL). Participant RD had 64-channel arrays in two regions of the hand knob in motor cortex, one positioned more medially (MCM) and one more laterally (MCL), as well as an array in the superior parietal lobule (SPL). An additional array targeting the supramarginal gyrus (SMG) in RD was excluded from all analyses, as it did not yield reliable task-related responses across sessions or experimental variations.

To investigate action encoding across intention and observation, we designed a task with a fully crossed 2 (hand: left/right) × 3 (action: lift, slide, rotate) × 2 (direction: left/right) structure (Fig. 1A). Direction refers to an abstract left/right factor, defined per action type (see Methods).In the intention condition, participants intended the cued action; in observation, they passively viewed a video of the same action. This structure allowed independent assessment of effector, action type, and direction encoding across formats.

**Figure 1:**
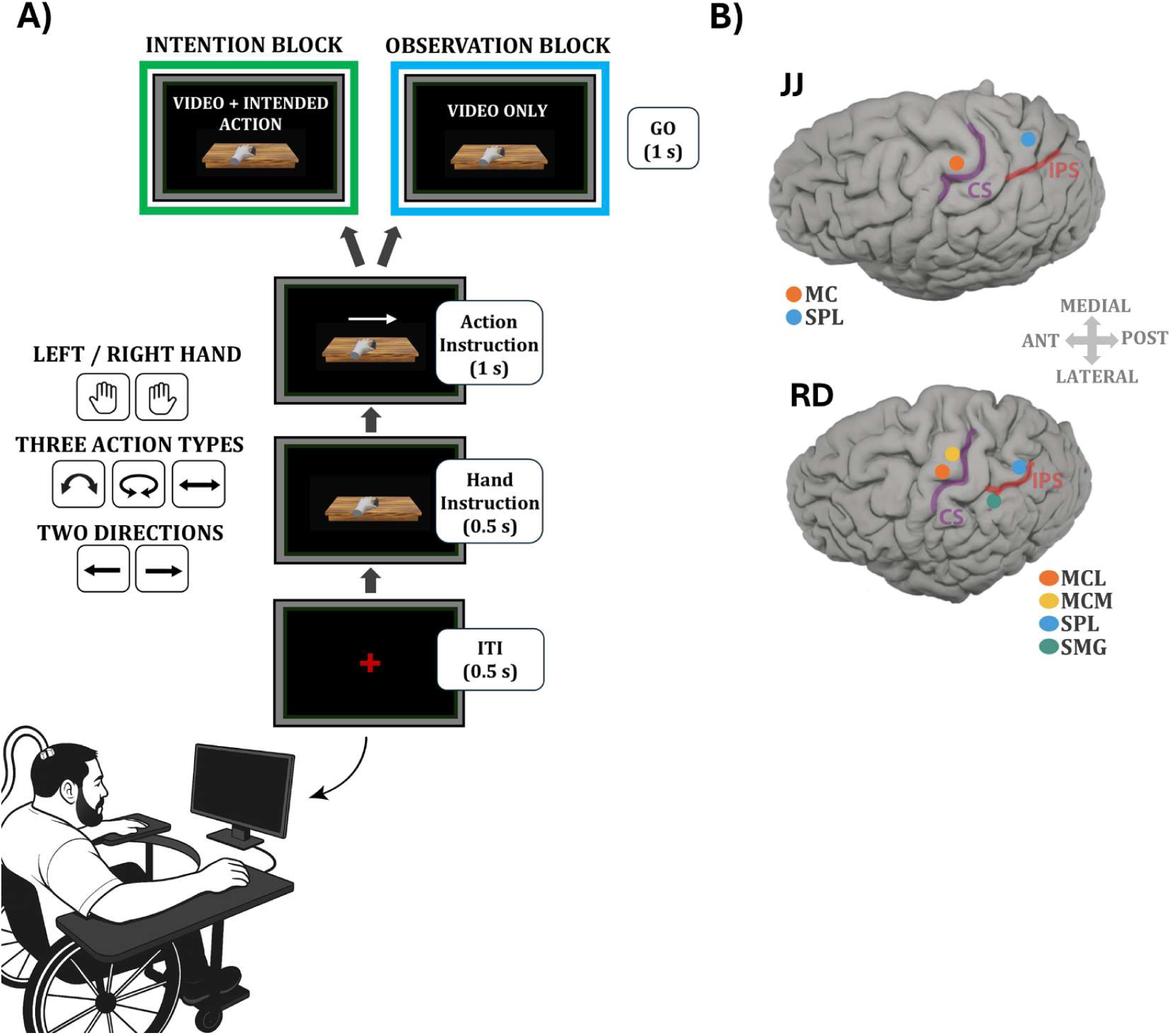
Experimental design and implant locations. **(A)** Action Intention and Observation Task. Each trial began with a 0.5 s inter-trial interval, followed by a 0.5 s hand cue and a 1 s symbolic action cue. The hand cue indicated the instructed effector (left or right), and the overlaid arrow specified the action type (slide, lift, rotate) and direction (leftward or rightward). The disappearance of the cue served as the go signal, followed by a 1s video of the corresponding action. Participants either intended (intention block) or passively viewed (observation block) the instructed movement. The design yielded 12 fully crossed conditions (2 hands × 3 actions × 2 directions). **(B)** Implant locations. Participant JJ had 96-channel arrays in motor cortex (MC) and superior parietal lobule (SPL). Participant RD had four 64-channel arrays: medial and lateral motor cortex (MCM, MCL) and posterior parietal cortex (SPL, SMG).

We recorded 421 units in MC and 326 in SPL from JJ, and 441 (MCM), 479 (MCL), and 532 (SPL) units from RD during the main task (Table S1). In RD, two further dissociation experiments yielded an additional 885 (MCM), 794 (MCL), and 1020 (SPL) units (Tables S2–S3), which were analyzed separately. Implant locations are shown in Figure 1B. Based on the quality factor classification (see Methods), 81 units in RD-MCM, 109 in RD-MCL, and 92 in RD-SPL were well isolated (quality 1–2), with the remainder classified as multiunit activity (quality 3–4). For JJ, whose arrays had been implanted for an extended period of time, the majority of signals were classified as multiunit activity, yielding only 5 well-isolated MC units and 4 well-isolated SPL units. For the dissociation tasks, the numbers of well-isolated units were 134 in RD-MCM, 107 in RD-MCL, and 254 in RD-SPL.

Eye-tracking confirmed stable fixation across all conditions. Figure S1 shows the two-dimensional distributions of gaze positions (in degrees of visual angle) aggregated across all trials. Across both participants and formats, gaze remained consistently centered, confirming compliance with fixation instructions throughout the experiment.

## Distinct tuning profiles in parietal and motor cortices across action formats

We first examined condition-specific responses at the single-unit level across regions. Figure 2A shows example units from RD: one in SPL tuned to action “lift”, and one in MCM tuned to right-hand actions. Both showed consistent tuning across formats. The SPL unit exhibited peak responses for lift at 2.1 s (159 spikes/s) during intention and at 2.0 s (82.5 spikes/s) during observation. The MCM unit showed sustained right-hand selectivity, peaking at 2.0 s (83.2 spikes/s) during intention and 1.7 s (48.5 spikes/s) during observation. However, consistent cross-format tuning was not ubiquitous; many neurons showed divergent selectivity across formats.

**Figure 2:**
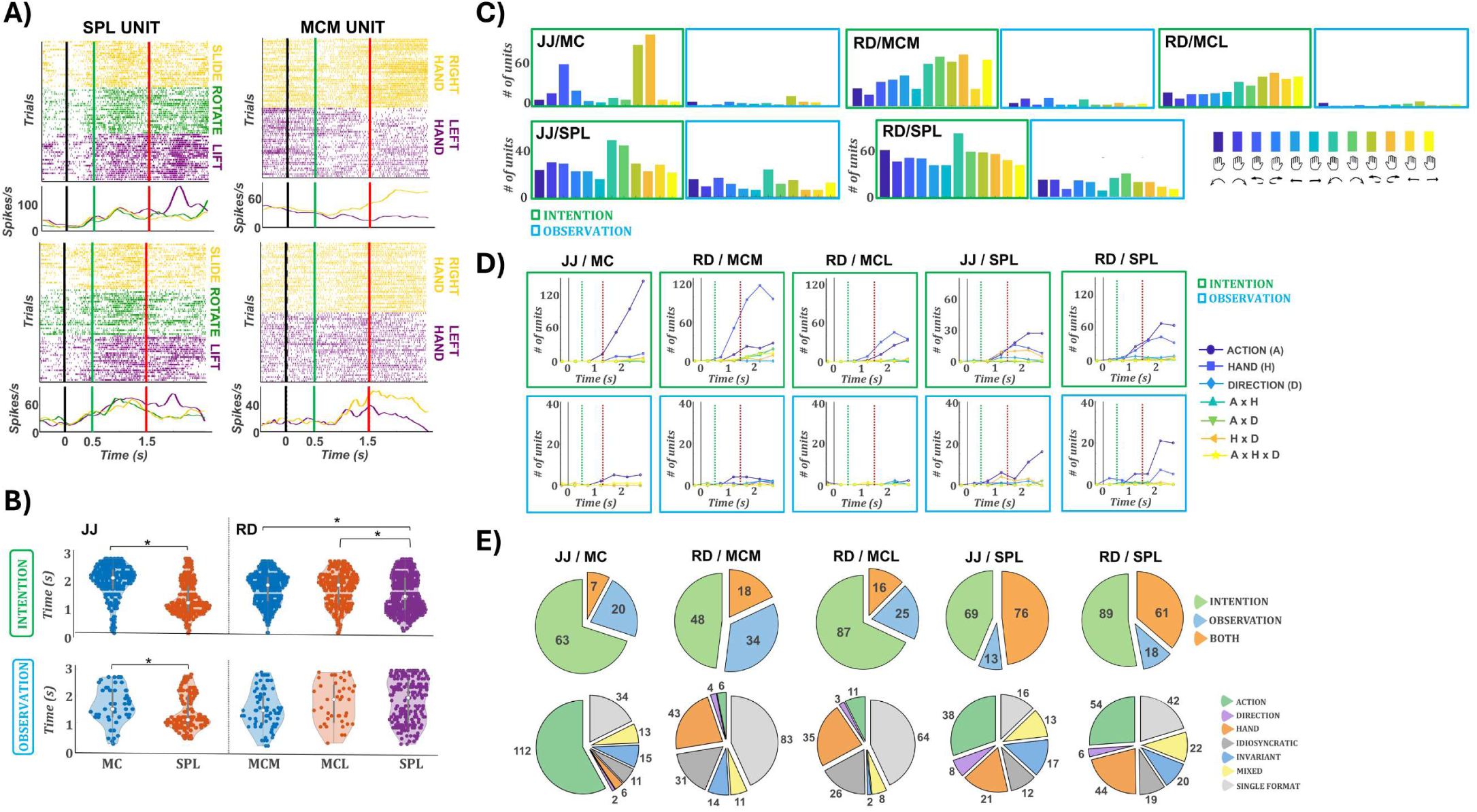
Single-unit selectivity across intention and observation. **(A)** Example neurons from RD in SPL (left) and MCM (right). Top: intention; bottom: observation. Trials are color-coded by action for SPL (yellow: slide, green: rotate, purple: lift) and by hand for MCM (yellow: right, purple: left). Below each raster, the corresponding PSTHs are shown using the same color scheme. Vertical lines indicate task events: the black line (time 0) marks the hand cue, the green line at 0.5 s the action cue, and the red line at 1.25 s the go cue. The SPL neuron exhibits clear selectivity for the lift action across both formats, while the MCM neuron responds selectively to the right hand. Both neurons show format-general responses. **(B)** Violin plots of onset latencies across all arrays for intention (top) and observation (bottom). Asterisks indicate statistically significant differences. In JJ, SPL units show significantly earlier responses than MC units in both formats. In RD, SPL responses are earlier than those in both MCM and MCL during intention, but not during observation. **(C)** Number of significantly responsive units per condition (Bonferroni-corrected t-test against baseline, ≥3 consecutive bins). Green and blue outlines correspond to intention and observation blocks, respectively. MC and MCM arrays show robust responses during intention but few during observation. In contrast, SPL arrays show substantial responses in both formats. **(D)** Three-way ANOVA tuning profiles over time (factors: action, hand, direction). Top: intention (green outline); bottom: observation (blue outline). Lines represent main effects and all two– and three-way interactions. In JJ, MC is strongly tuned to action. MCM and MCL (RD) are predominantly tuned to hand, followed by action. SPL in both participants shows dominant action tuning and secondary hand tuning. Interaction terms are rare. During observation, only SPL arrays preserve action tuning. (**E)** Format selectivity (top row): number of units with significant responses in intention only (green), observation only (blue), or both (orange). These counts reflect response presence but not tuning congruency. Linear model classification (bottom row): units are assigned to their best-fitting model based on cross-validated R². Categories include shared action (green), shared hand (orange), shared direction (purple), invariant (blue), mixed (yellow), idiosyncratic (dark gray), and single-format (light gray). Unselective units are not shown. SPL arrays in both participants contain the fewest single-format units and the highest proportion of shared action models.

To compare temporal dynamics, we computed response latencies per condition and unit, defined as the center of the first of three consecutive bins (100 ms each) significantly above baseline. Figure 2B shows latency distributions across regions and formats. During intention, SPL responses preceded those in motor cortex (median latencies: JJ/SPL: 1.2 s; RD/SPL: 1.4 s; MC/MCM/MCL: 1.9–2.1 s; ANOVA p < 0.001). With the effector cue at 0 s, the action and direction cue at 0.5 s, and the go cue at 1.5 s, these latencies indicate that SPL activity emerged ∼700–900 ms after the action cue and ∼100–300 ms before the go cue, whereas MC responses appeared ∼400–600 ms after the go cue. During observation, SPL also preceded MC in JJ (1.1 vs. 1.5 s, p = 0.02), but no significant timing differences were found in RD. These results point to a leading role of SPL during action intention, with less consistent dynamics during observation.

We quantified the number of units responsive to at least one condition in each format and brain area. A unit was considered responsive if it showed three consecutive time bins significantly above baseline (see Methods). Figure 2C shows the distribution of responsive units per condition, separately for intention (top row, green) and observation (bottom row, blue). In motor cortex, substantially more units responded during intention than observation: JJ/MC: 220 vs. 50; RD/MCM: 215 vs. 40; RD/MCL: 172 vs. 39. Notably, in JJ’s MC, responses were strongest for right-hand rotation, whereas MCM and MCL exhibited broader tuning, though with a right-hand bias. In SPL, responses were widespread across both formats. During intention, 139 (JJ) and 256 (RD) units were responsive; during observation, 88(JJ) and 137(RD). This exceeds the number observed in any motor cortex array, indicating that SPL supports robust responsiveness across formats, whereas motor cortex is primarily active during intention.

To assess tuning to specific task features, we performed a three-way ANOVA on binned firing rates with action type, effector (hand), and direction as factors (Fig. 2D). In motor cortex during intention, tuning patterns differed by participant. In JJ’s MC, action tuning dominated (peak: 144 units at 2.75 s), while effector tuning was minimal (14 units). In RD, MCM showed strong effector tuning (118 units at 2.25 s), with weaker action (28) and interaction tuning (e.g., action×hand: 18). MCL showed tuning for both hand (45) and action (33), with hand remaining dominant. SPL exhibited a distinct profile. In both participants, action was the most commonly tuned feature during intention (JJ: 27; RD: 58), followed by effector (JJ: 16; RD: 37). This structure was largely preserved during observation (JJ: 16; RD: 21 action-tuned units), whereas observation-related tuning in motor cortex was minimal (≤5 units for all features in RD; 5 action-tuned units in JJ’s MC). Across regions, effector tuning tended to emerge earlier than action, particularly during intention suggesting a sequential encoding of task parameters. For example, hand tuning peaked around 2.25 s in MCM and MCL, while action peaked at 2.75 s. SPL showed a similar temporal shift. Interaction terms were rarely significant, suggesting that most units were modulated by single task features. Overall, motor cortex was dominated by effector or action tuning during intention, while SPL consistently encoded action type in both formats.

We next quantified format-specific responsiveness by counting units active for at least one condition during intention only, observation only, or both (Fig. 2E, top row). In motor cortex, most units were responsive exclusively during intention. In JJ’s MC, 63 units were intention-only, compared to 20 observation-only and 7 responsive in both formats. Similar distributions were found in RD: MCM (48 intention-only, 34 observation-only, 18 both) and MCL (87, 25, and 16, respectively). In contrast, SPL showed a higher proportion of units responsive across both formats. In total, 76 units in JJ and 61 in RD responded in both formats, nearly matching or exceeding the format-specific counts. This suggests that SPL encodes actions in a more format-invariant manner than motor cortex.

To characterize tuning structure and generalization, we applied a linear model framework adapted from Chivukula et al. 2025 ^22^. For each unit, we fit a full factorial model using action, hand, and direction (including all interactions), separately for intention and observation. Units were assigned to the best-fitting model via stratified 5-fold cross-validation. Shared tuning categories (consistent tuning to a given feature across both intention and observation formats) included action, hand, direction, mixed (e.g., action ×hand), or invariant (consistent across all 12 conditions). Units were labeled idiosyncratic if they were tuned in both formats but with unrelated profiles, single-format if tuned in only one, and unselective if no model exceeded R² > 0.01 or reached significance after FDR correction. Figure 2E (bottom) shows the distribution of model classifications across regions. In RD, MCM and MCL were dominated by single-format units (83 and 64), followed by shared hand (43 and 35), consistent with effector-dominant tuning (Fig. 2D). JJ’s MC showed a strikingly different pattern: despite few observation-responsive units (Fig. 2E, top), the majority were best fit by a shared action model (112), with only 34 classified as single-format. SPL in both participants exhibited consistent format-general structure. Shared action was the most frequent model (JJ: 38; RD: 54), followed by shared hand (21; 44), invariant (17; 20), and mixed models (13; 22). Single-format tuning was also present, particularly in RD (42 units). These findings further support the view that SPL contains a population of units that encode action features in a consistent and generalizable format across observation and intention.

To ensure that collapsing leftward and rightward variants of each action did not inflate shared-action fits, we repeated the model comparison using a 6-level action factor (action × direction; Fig. S2). The overall distribution of model fits remained consistent. Across areas, the 3-action model continued to dominate, particularly in PPC (JJ SPL: 52 action(3) vs 25 action(6) units; RD SPL: 29 action(3) vs 12 action(6)). In motor cortex, a similar profile was observed (JJ MC: 96 action(3) vs 28 action(6); RD MCM: 8 action(3) vs 6 action(6), RD MCL: 9 action(3) vs 5 action(6)). Thus, redefining actions as direction-specific variants revealed only a minor subset of additional tuned units, confirming that the dominance of action(3) tuning (particularly in PPC) reflects genuine encoding of action identity rather than conflation of directional movements.

At the single unit level SPL exhibited robust, format-general tuning across participants. In both JJ and RD, many units responded in both formats (76 and 61, respectively; Fig. 2C–E), with action tuning aligned in time and shared action emerging as the most frequent model. In contrast, motor cortex responses were predominantly intention specific. JJ’s MC showed strong action tuning, and RD’s MCM and MCL were dominated by effector tuning, but observation-related responses were sparse. Most MC units were classified as single-format, except in JJ, where shared action tuning was prevalent despite limited observation responsiveness. This apparent discrepancy reflects the difference between the baseline test, which detects only strong increases in firing, and the model analysis, which can reveal shared tuning structure even when observation responses are weak. Direction tuning was negligible across all regions. Overall, SPL supported consistent, format-invariant encoding, while motor cortex tuning was more format-dependent.

## Cross-format representational similarity is robust in SPL but conditional or absent in MC

Single-unit analyses revealed strong format independent tuning in SPL and single format, intention-dominant responses in motor cortex. We then examined whether this representational structure was preserved across formats at the population level. To test this, we applied representational similarity analysis (RSA) to quantify condition-specific similarities in population activity within and across formats (see Methods). We included only task-relevant units, as defined by the linear model analysis (see Methods), with the following counts per array: MC/JJ: 163, SPL/JJ: 101, MCM/RD: 117, MCL/RD: 88, and SPL/RD: 165.

Figure 3A shows cross-format RSA matrices for each array and task variable (action, hand, direction). Corresponding within-format matrices are shown in Figure S3, and analysis using different time windows and including all recorded units are presented in Figure S4. In JJ’s MC, cross-format similarity was specific to rotation (R² = 0.31); lift and slide were near zero. In RD’s MCM, correlations were strong for the right hand (0.71) and moderate for lift (0.53) and rotation (0.52); left-hand and slide were weaker (≤0.24). RD’s MCL showed no meaningful correlations (R² < 0.12). These patterns reflect selective generalization in MC/JJ and MCM/RD, aligned with their dominant intention-driven features (rotation and right-hand tuning; Fig. 2C–D), despite minimal observation tuning at the unit level. In contrast, SPL showed robust, distributed cross-format similarity across all task variables in both participants. Correlations were high for all actions (JJ: 0.59–0.72; RD: 0.71–0.79), hands (JJ: 0.63–0.68; RD: 0.76–0.78), and directions (JJ: 0.63–0.64; RD: 0.73–0.79), consistent with stable representational structure.

**Figure 3:**
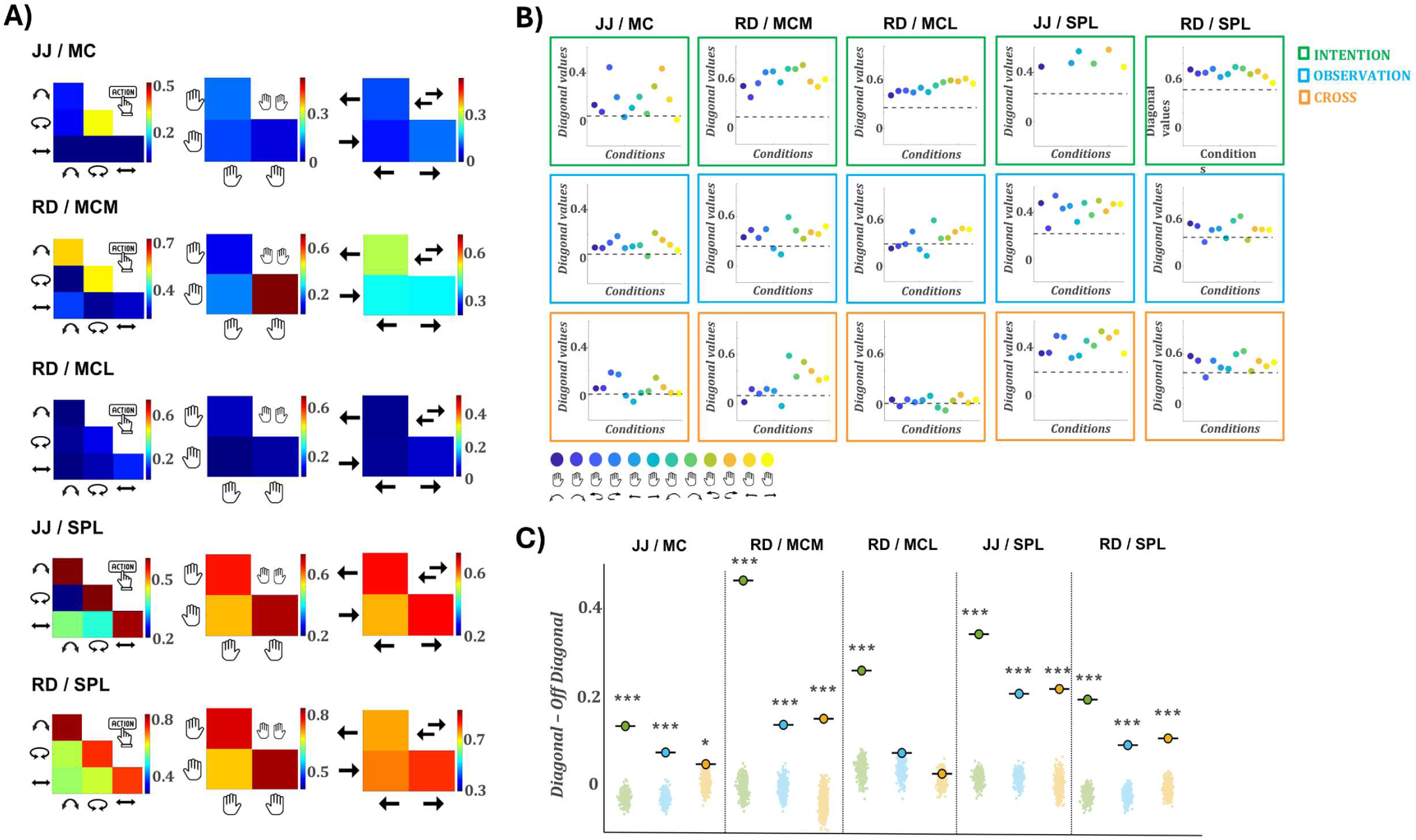
Representational similarity analysis (RSA) within and between formats. **(A)** Cross-format RSA heatmaps for each array, computed separately for action (left column), hand (middle), and direction (right). Each matrix shows the correlation between condition-specific population vectors from intention and observation formats. MC (JJ) and MCM (RD) show elevated cross-format correlations for specific features only, rotation and right hand, respectively. MCL shows no cross-format similarity. SPL arrays in both participants exhibit clear diagonal structure across all three variables, indicating preserved population representations between intention and observation. (**B)** Diagonal values of 12-condition RSA matrices for each array and format. Top row: within-format intention; middle: within-format observation; bottom: cross-format. Each dot represents one of the 12 conditions; dashed lines indicate the average off-diagonal value. Within-format diagonals are consistently above the off-diagonal mean, confirming meaningful condition structure. Cross-format results reveal isolated high values in JJ/MC (rotation) and RD/MCM (right hand), matching the heatmaps in A. In MCL, all diagonal values are near the off-diagonal mean, confirming a lack of structure. SPL arrays again show consistently elevated diagonals in both participants, indicating generalizable encoding across formats. **(C)** Permutation test results for each array and format. Each panel shows the real diagonal–off-diagonal difference (bold marker) against a null distribution from label-shuffied permutations (transparent dots). Colors indicate format: green for intention, blue for observation, orange for cross. Asterisks denote significance compared to the null distribution (*p < 0.05; ***p < 0.001). In MCL, only intention responses differ significantly from the null.

Figure 3B shows RSA diagonal values for the 12 task conditions in intention, observation, and cross-format analyses. During intention, diagonal values consistently exceeded the off-diagonal mean across all arrays, reflecting strong condition-specific encoding, most pronounced in SPL, MCL, and MCM, and weaker in MC/JJ. Observation responses showed similar but reduced structure across arrays. Cross-format RSA revealed selective structure in MC/JJ and MCM/RD. In MC/JJ, rotation actions showed elevated diagonal values and in MCM/RD, the highest values corresponded to right-hand actions, consistent with effector-specific generalization. MCL showed no meaningful cross-format structure, with all diagonals near the baseline. In contrast, SPL in both participants showed consistently elevated diagonal values across nearly all conditions, reflecting robust and generalizable cross-format representations.

Lastly, to quantify structure, we compared the difference between mean diagonal and off-diagonal RSA values against a null distribution generated via 1,000 label permutations (Fig. 3C). In SPL, all formats showed significant structure in both participants (JJ: int = 0.35, obs = 0.21, cross = 0.22; RD: int = 0.20, obs = 0.10, cross = 0.10; all p < 0.001). MC/JJ and MCM/RD also showed significant structure in all formats (MC: 0.14, 0.08, 0.05; MCM: 0.47, 0.14, 0.15; all p < 0.05). MCL/RD reached significance only during intention (0.25, p < 0.001); observation and cross-format effects were nonsignificant (p > 0.05).

Together, these results demonstrate that SPL consistently encodes task structure across formats at the population level, while in motor cortex, representational similarity is either absent or limited to specific, strongly encoded features, highlighting the importance of population-level analyses in revealing structure that may not be apparent from single-unit responses alone.

## SPL supports robust cross-format decoding, while motor cortex shows asymmetric or absent generalization

We assessed how reliably task features could be extracted from population activity over time, by performing time-resolved decoding analyses for action type, effector, and movement direction, separately for intention (Figure 4A) and observation (Figure 4B) trials. Figure S5 displays decoding accuracy and confusion matrices for all 12 task conditions. During intention all arrays showed robust decoding of action type and effector. Action decoding peaked above 90% in SPL for both participants (JJ: 91.1%; RD: 95.8%) and ranged from ∼74–83% in the motor cortex arrays (JJ/MC: 73.6%; RD/MCM: 77.9%, MCL: 83.2%). Effector decoding was similarly strong, peaking above 93% in all of RD’s arrays (MCM: 93.7%, MCL: 93.8%, SPL: 93.0%), and above 75% in JJ (MC: 76.4%, SPL: 86.9%). Direction decoding remained weak across regions. Effector decoding consistently peaked earlier than action, supporting a sequential encoding scheme. During observation, decoding was strongest in SPL (action: JJ: 77.7%, RD: 77.1%), with moderate effector (JJ: 65.7%, RD: 67.4%) and direction decoding. Motor cortex decoding during observation was weak, except for action in MC/JJ (72.2%). Thus, SPL reliably encoded action type across formats. Across sessions (Figure S6), decoding performance remained stable: action and hand decoding were consistently above chance for intention in all regions, with SPL showing the highest action decoding during observation, while direction decoding remained uniformly poor.

**Figure 4:**
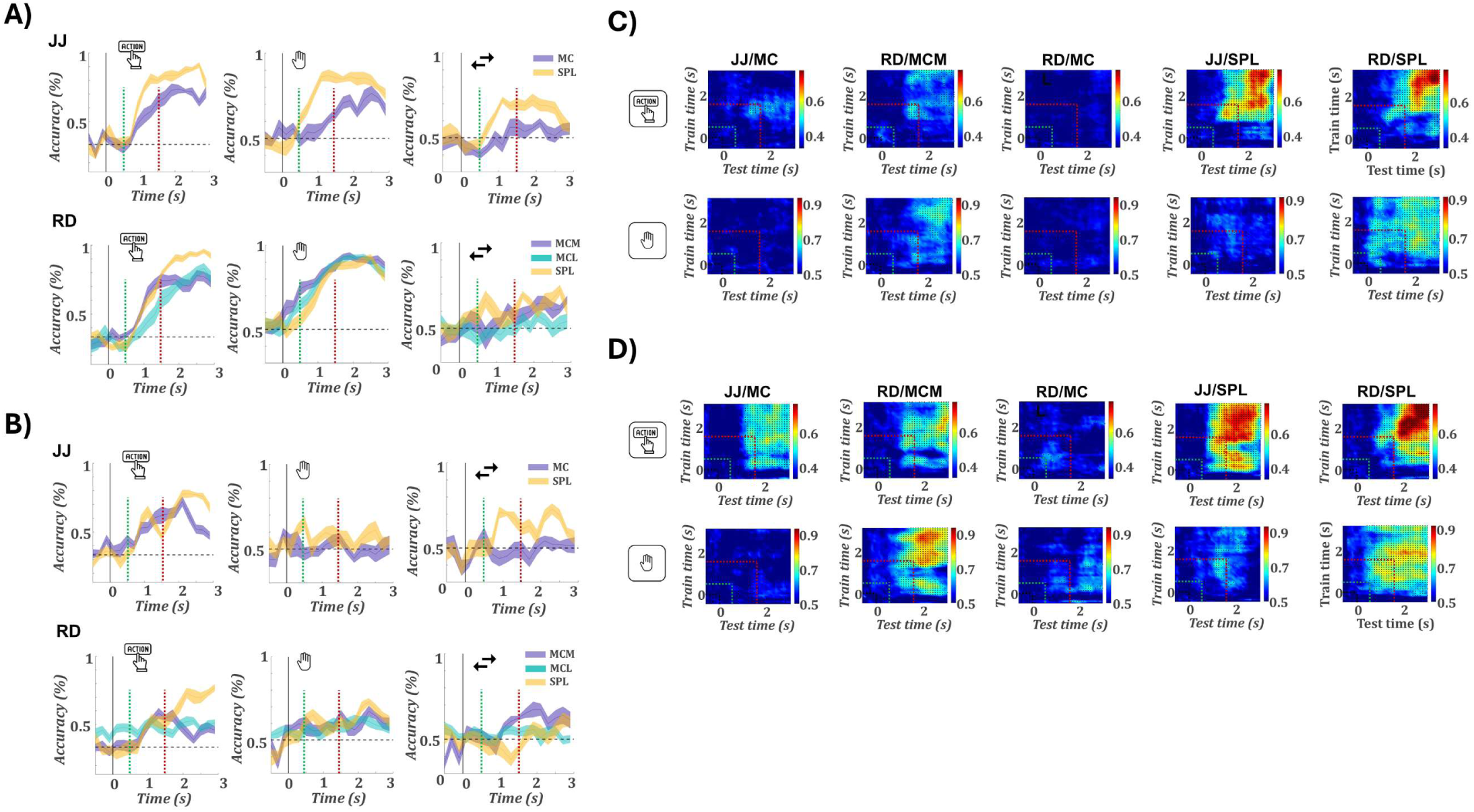
Decoding task variables within and across formats. **(A)** Time-resolved decoding accuracy for action type (left), hand (middle), and direction (right) during intention. Top row: JJ (MC, SPL); bottom row: RD (MCM, MCL, SPL). All arrays decode action and hand well above chance. Vertical lines mark the action cue (green) and go cue (red). Shaded regions denote SEM across cross validations. The dashed horizontal line marks chance level (**B)** Same analysis for the observation format. Decoding performance is generally reduced compared to intention. SPL continues to show robust decoding, particularly for action type. **(C)** Cross-format decoding: classifiers trained on intention data and tested on observation. Each heatmap shows decoding accuracy across all combinations of training (y-axis) and testing (x-axis) time bins. Green and red dashed lines indicate the onset of the symbolic action cue and the go cue, respectively. Significant decoding time points (permutation test, p<0.05) are marked with dots. SPL exhibits strong generalization of action representations in both participants. MCL shows no evidence of generalization. **(D)** Reverse cross-format decoding: classifiers trained on observation data and tested on intention. SPL again supports robust generalization for action type and, to a lesser extent, hand. RD/MCM shows strong decoding for action and weaker generalization for hand, while JJ/MC also decodes action. MCL does not support generalization in either direction.

To assess generalization across formats, we performed cross-temporal decoding by training classifiers on one format and testing on the other, across all time point pairs (Figs. 4C–D). Analyses were run bidirectionally: Intention → Observation (Fig. 4C) and Observation → Intention (Fig. 4D), for action and hand. Results for movement direction are shown in Figure S7. Decoding accuracy matrices were statistically thresholded via permutation testing (see Methods). In the Intention → Observation direction, SPL showed robust cross-decoding of action in both participants (JJ: 77.1% at 2.7→2.6 s; RD: 79.9% at 2.6→2.8 s), with strong hand decoding in JJ (78.5% at 1.1→2.1 s). A small number of additional significant effects were also observed in MC of JJ and MCM of RD, with the strongest in MCM for hand (72.2% at 2.1→2.5 s). No significant generalization was detected in MCL. In the Observation → Intention direction (Fig. 4D), SPL exhibited robust generalization for both action (JJ: 77.0% at 2.5→2.5 s; RD: 92.4% at 2.9→2.8 s) and hand (RD: 86.1% at 1.8→2.1 s; JJ: 71.5% at 1.2→1.2 s), with tightly aligned timing across formats. Interestingly, motor cortex also showed significant decoding: MC/JJ generalized action (62.5% at 1.5→2.1 s), and MCM/RD generalized both hand (88.2% at 2.5→1.9 s) and action (66.7% at 2.2→2.8 s). No effects were found in MCL. The unidirectional decoding observed in MC and MCM is particularly notable given the weak or absent within-format decoding during observation in these regions. This suggests that, even in the absence of overt task selectivity, observation-related activity may retain structured components aligned with intention representations. In contrast, the robust and bidirectional decoding in SPL indicates the presence of a stable, format-invariant code, particularly for action type, consistent with its generalized encoding across tasks and formats.

## SPL and motor cortex exhibit distinct representational geometries across action formats

A central question in understanding action encoding is whether population activity occupies similar geometric structure across cognitive states. To address this, we examined the organization of neural trajectories during intention and observation. Trial-averaged responses were projected into a shared PCA space based on condition means (Fig. 5A; Fig. S8), providing a geometric perspective on population structure. These trajectories provide a striking and intuitive geometric perspective on the population structure underlying action encoding. Table S2 reports the variance explained by the first three principal components for intention, observation, and the combined dataset, separately for each array and task variable. These values confirm that the projections capture sufficient variance to support meaningful trajectory analysis, with total explained variance exceeding 60% in all cases. They also highlight format-specific differences: observation variance was consistently lower than intention in motor cortex, while SPL showed comparable variance across formats. In MC of JJ, trajectories appeared similar across formats only for the rotation action, consistent with its selective generalization in RSA (Fig. 3A) and decoding (Fig. 4D). In MCM of RD, responses were more similar for right-hand conditions, in line with effector-specific cross-format structure. MCL showed clearly segregated trajectories across formats for all conditions, matching the absence of generalization observed throughout prior analyses. In contrast, SPL exhibited qualitatively similar trajectories for all actions across formats, consistent with its robust and generalized encoding. To quantify cross-format similarity, we applied Procrustes analysis between observation and intention trajectories for each task variable and condition (Fig. 5B–C). This method estimates the best-fitting linear transformation (translation, rotation, scaling) to align observation trajectories onto their intention counterparts and returns a distance metric reflecting residual dissimilarity. SPL showed low alignment distances across all conditions (d < 0.1), indicating consistent geometric overlap. In MC of JJ, low distance was found only for rotation (d = 0.25), with poor alignment for lift (0.89) and slide (0.64). In MCM of RD, right-hand trajectories aligned more closely (d = 0.13) than left-hand ones (0.30). No condition in MCL yielded meaningful alignment. These results confirm that SPL supports a shared representational geometry across formats, while MC and MCM exhibit selective overlap, and MCL none. Although Procrustes alignment does not mean that observation and intention trajectories occupy the same neural space, the transformation removes differences in translation, rotation, and scale to reveal their intrinsic geometry. Successful alignment therefore indicates that observation and intention preserve a similar representational structure that can be linearly mapped across formats despite global shifts in activity or response gain.

**Figure 5:**
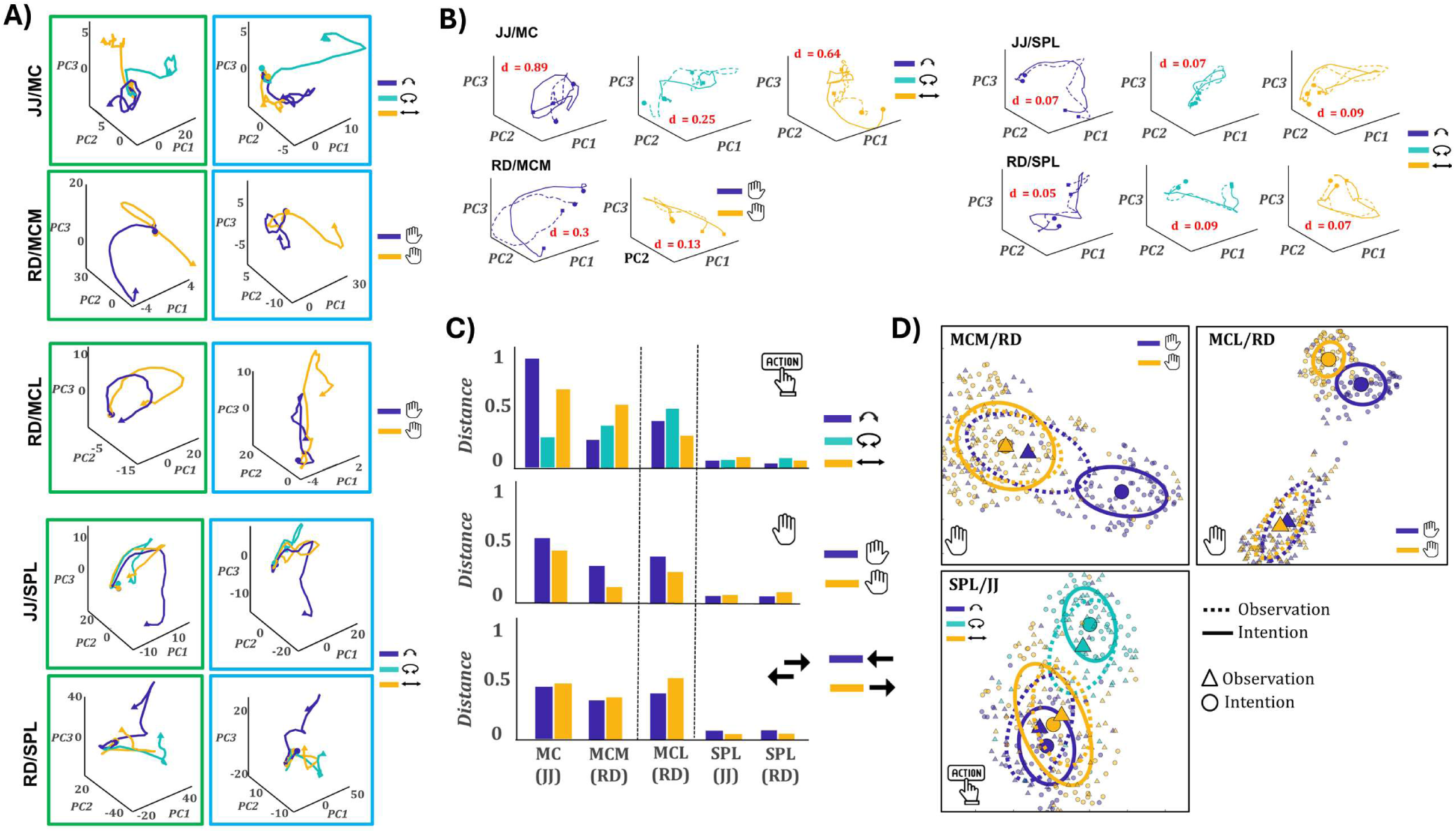
Representational geometry of neural representations across formats. **A)** Example PCA trajectories for intention (left column) and observation (right column, blue outline), plotted in the first three principal components for different task variables: action (JJ/MC, JJ/SPL, RD/SPL) and hand (RD/MCL, RD/MCM). In JJ/MC and RD/MCM, single trajectories (rotation and right hand, respectively) show similar geometry across formats, while other trajectories diverge. In RD/MCL, trajectories are completely dissociated between formats. In contrast, in SPL (both JJ and RD), all action trajectories appear nearly identical across formats. **B)** Procrustes-aligned trajectories from the same examples in A, with intention shown as solid lines and observation as dashed lines. **C)** Distances (**d**) quantify dissimilarity after alignment. In JJ/MC and RD/MCM, only one specific condition (rotation and right hand, respectively) aligns well across formats (d < 0.3). In SPL (JJ and RD), all action trajectories align tightly with d < 0.1, indicating nearly identical representational geometry across formats. **C)** Summary of alignment distances for all arrays and task variables (action, hand, direction). SPL clearly stands out, with alignment distances below 0.2 across all task features, in contrast to MC and MCM where only selective geometries generalize, and MCL where alignment fails entirely. **D)** UMAP embeddings illustrating three representative cases of cross-format geometry. Triangles represent observation trials and circles represent intention. Colors denote task conditions, and ellipses enclose clusters (solid: intention, dashed: observation). Top left (RD/MCM, hand): Right-hand trials from both formats cluster together, while left-hand trials form separate clusters, indicating conditional overlap. Top right (RD/MCL, hand): Intention and observation clusters are completely segregated, indicating no shared geometry. Bottom (JJ/SPL, action): Clusters for all three actions overlap across formats, indicating a fully shared representational geometry.

To further assess population structure across formats, we applied Uniform Manifold Approximation and Projection (UMAP) to embed single-trial activity from both intention and observation into a shared low-dimensional space (Fig. 5D; Fig. S9). Trials were color-coded by condition, and ellipses were fit to the format-specific distributions to visualize condition clustering and cross-format overlap. Figure 5D presents three representative UMAP embeddings (MCM/RD (effector), MCL/RD (effector), and SPL/JJ (action type). Three distinct regimes emerged. In MCM of RD, responses for right-hand trials formed overlapping clusters across formats, whereas left-hand trials remained separated, mirroring the effector-specific generalization seen in RSA, decoding, and PCA. In MCL, intention and observation trials occupied distinct, non-overlapping regions, consistent with the complete absence of cross-format generalization. In contrast, SPL of JJ showed strong cross-format overlap for all action types, with well-separated but aligned clusters, particularly for rotation. These embeddings reinforce the dissociation across regions: SPL supports robust, format-general population codes; MCM exhibits conditional overlap; and MCL remains format-specific.

## High-gamma LFP responses reveal format-consistent population encoding at the single channel level

To evaluate neural population structure beyond spiking activity, we analyzed high-gamma (60–120 Hz) LFP power as a complementary population-level metric (Fig. 6). Across all arrays and participants, all channels showed significant modulation for at least one condition in both intention and observation. Strikingly, all motor cortex arrays exhibited robust observation-related responses, in contrast to the sparse or absent observation tuning in the SUA/MUA data. Figure 6A illustrates these effects via time–frequency plots, showing consistent condition-specific activations across formats e.g., rotation tuning in JJ’s MC, and right-hand selectivity in MCM and MCL of RD. Figure 6B quantifies the distribution of significantly modulated channels across conditions, highlighting that format-specific tuning was preserved: rotation tuning in JJ MC and effector tuning in MCM/MCL of RD appeared in both formats. SPL again exhibited widespread modulation across all conditions in both participants. Figure 6C shows the evolution of tuning over time. In motor cortex, tuning profiles were nearly identical between intention and observation: action-selectivity in JJ’s MC and hand-selectivity in MCM and MCL of RD. SPL showed consistent tuning for action during observation in both participants; during intention, tuning was action-dominant in JJ and hand-dominant in RD. Decoding from LFPs was weaker than from SUA but followed similar trends (Fig. 6D, S10). During intention, both action and hand could be decoded above chance in all arrays (action: JJ/MC: 57.6%, SPL: 70.4%; RD/MCM: 47.8%, MCL: 47.4%, SPL: 60.8%; hand: JJ/MC: 63.3%, SPL: 70.7%; RD/MCM: 73.2%, MCL: 64.4%, SPL: 67.7%).

**Figure 6:**
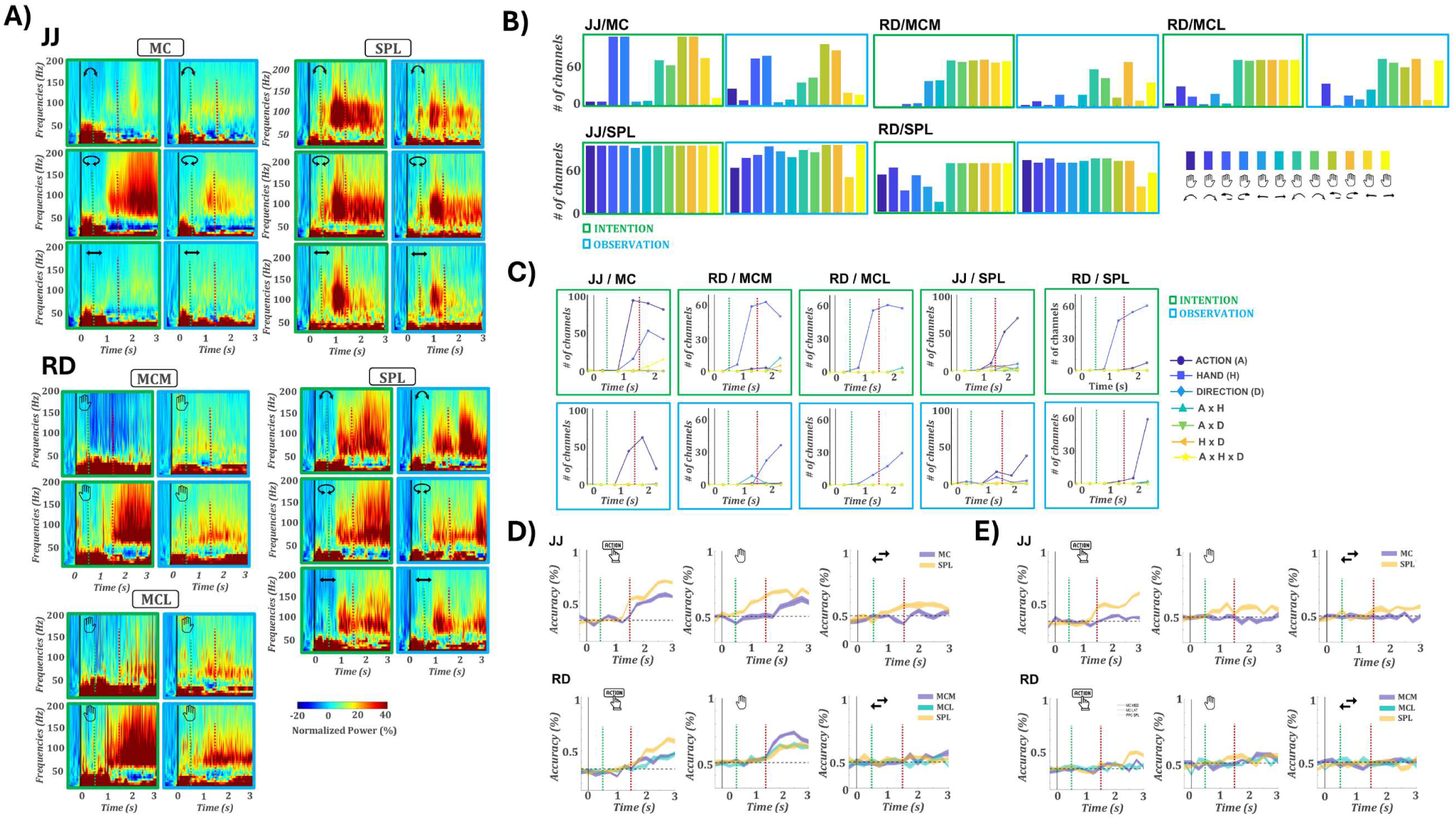
High-frequency LFP activity across formats and cortical regions. **(A)** Example spectrograms showing event-related changes in LFP power for selected conditions: action (JJ/MC, JJ/SPL, RD/SPL) and hand (RD/MCM, RD/MCL). Colors indicate percentage power change relative to baseline. Within each array, the left column (green outline) shows intention and the right column (blue outline) shows observation. In MC arrays, gamma-band activity is highly selective: e.g., rotation in JJ and right-hand responses in RD. Notably, similar selectivity patterns are also visible during observation. SPL arrays respond broadly to all actions in both formats. **(B)** Number of channels with significant power increases (t-test vs. baseline, Bonferroni-corrected, *p* < 0.05 for ≥3 consecutive 100 ms bins). Green and blue outlines correspond to intention and observation, respectively. As seen in the spectrograms, MC/MCM arrays show selectivity for specific conditions (rotation in JJ/MC, right hand in RD/MCM) during intention, which is preserved during observation. SPL channels exhibit widespread activation across both formats and multiple conditions. **(C)** Three-way ANOVA tuning profiles over time (factors: action, hand, direction). Top: intention (green outline); bottom: observation (blue outline). Lines indicate the number of channels significantly tuned to each main effect and interaction. MC/MCM arrays show clear tuning for specific task variables (e.g., action in JJ, hand in RD) during intention, and these patterns are preserved during observation. SPL shows broad tuning to action (JJ) and hand (RD) during intention, and action type during observation. **(D)** Within-format decoding of action, hand, and direction during intention. Top: JJ; bottom: RD. Each line shows the average decoding accuracy across cross-validation folds; shaded regions indicate ±SEM. The dashed horizontal line marks chance level. Vertical green and red dashed lines indicate the onset of the symbolic action cue and the go cue, respectively. All arrays show above-chance decoding for action and hand, though peak accuracy remains modest (<75%). **(E)** Same as D, for observation. Only SPL shows above-chance decoding, limited to action type in JJ and weakly in RD. Other regions fail to decode task features reliably during observation.

During observation, only SPL supported above-chance decoding for action (JJ: 60.1%; RD: 48.9%). Cross-format decoding (Fig. S11) recapitulated key spiking results: SPL showed robust bidirectional generalization; MCM exhibited unidirectional hand generalization; and MC of JJ showed a weak action effect. These findings indicate that high-gamma activity captures structured population level encoding even when single-unit selectivity appears sparse. This pattern should not be interpreted as evidence for a distinct representational format in the LFPs. Rather, the discrepancy reflects differences in sampling: our spike recordings capture a limited subset of neurons, whereas high-gamma signals pool over a much broader local population. Consistent with this view, spike-based population analyses (RSA, cross-decoding, trajectory analyses) already uncovered latent representational geometry despite weak tuning at the single-unit level. High-gamma activity expressed this geometry more robustly, underscoring how population-level signals can expose consistent representational structure that is only partially evident in sparsely sampled units. In SPL, the convergence of SUA, LFP, and population metrics reinforces its role as a stable, format-general hub for action representation.

## SPL Flexibly Represents Observed Actions Only When They Are Behaviorally Relevant

To determine whether neural representations of instructed and observed actions are encoded in parallel or selectively modulated by task demands, we designed two dissociation tasks that explicitly separated intention from observation (Fig. 7A). On every trial, participant RD was presented with both an instruction cue (specifying which hand and action to perform) and a concurrent video (showing a hand performing an action). The instruction and video could be either congruent or incongruent, but task demands determined which source of information was behaviorally relevant. In the no-probe variant, task blocks defined the relevant source: during the intention block, RD executed the instructed action while the concurrent video was present but not relevant to the task; during the observation block, RD passively viewed the video while the instruction cue was present but not relevant. In the probe variant, RD again executed the instructed action while a video played, but after movement completion was required to report a feature of the video (action or hand) with a saccade, making both the instruction and the video behaviorally relevant. This design enabled us to dissociate intention-related and observation-related activity and to test how behavioral relevance affects neural representations across regions. Session-level unit counts for both experiments are reported in Tables S3, S4. In the no-probe intention block, all areas encoded the instructed action, with strong hand tuning in MCM (163 units at 1.75 s) and MCL (90 at 1.25 s), and stronger action tuning in SPL (60 units at 2.25 s) (Fig. 7B, green). However, none of the areas encoded the concurrent video action while intention was underway. In the no-probe observation block, where no movement was intended, tuning for the instructed (now irrelevant) action dropped to baseline in all areas, while SPL selectively represented features of the video, with 23 units tuned at 2.25, predominantly to action identity rather than hand or interaction terms (Fig. 7B, blue). In the probe variant, tuning for the instructed action remained strong in motor cortex (MCM: 122 hand-tuned units at 2.25 s; MCL: 74 at 1.25 s) and SPL (96 action-tuned at 1.25 s) (Fig 7B, gray).Crucially, SPL exhibited robust selectivity for the video features (29 units at 2.25 s), in sharp contrast to the no-probe variant where the same visual input was present but elicited no selectivity. Motor cortex, by comparison, again showed <5 tuned units for the video features (Fig. 7b, gray).

**Figure 7:**
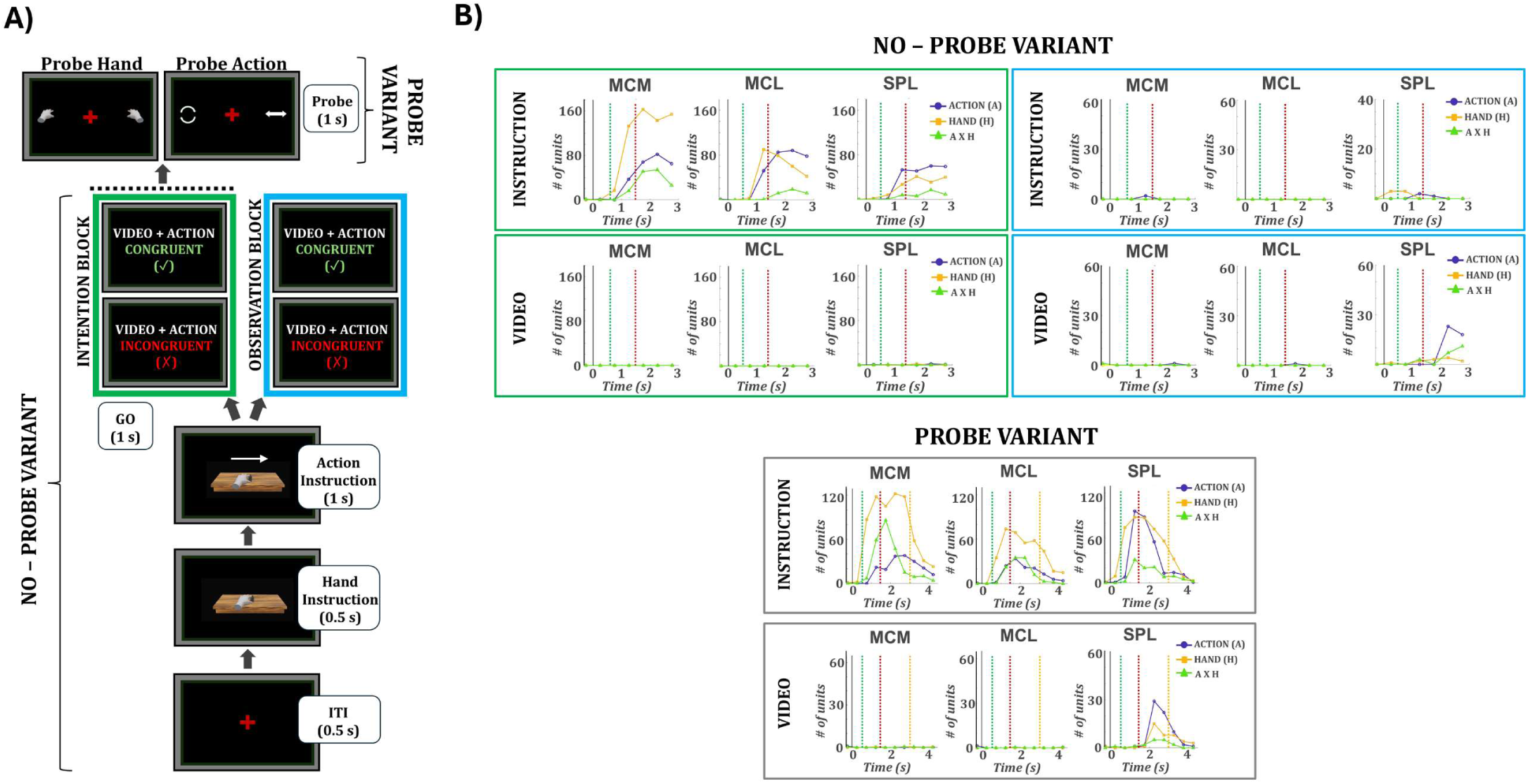
Dissociation tasks: neural tuning for instructed and observed actions. **(A)** Experimental design. Each trial began with hand and action instructions, followed by a video that could be congruent or incongruent with the instructed action. In the probe variant, a post-trial cue required a saccade to report either the video action or the video hand. In the no-probe variant, participants executed the instructed action while ignoring the video (intention block) or passively viewed the video while ignoring the instruction (observation block). **(B)** Time-resolved tuning. Green outline: no probe variant – instruction block; blue outline: no probe variant – observation block; gray outline: probe variant. Each plot shows the number of units tuned to the instructed (top row) or video (bottom row) action, hand, or their interaction in MCM, MCL, and SPL.

Decoding results mirrored the tuning patterns (Fig. 8A). During the no-probe intention block, instructed actions were decoded with high accuracy across all regions (peak accuracy: MCM: 99.4%, MCL: 97.5% SPL: 91.9%; Fig. 8A, green), but decoding of the concurrent video action remained near chance (<37% in all areas). In the observation block, decoding of the instructed (irrelevant) action dropped below 42% in all regions, while decoding of the video action rose sharply in SPL (66.3% at 2.5 s) but remained weak in motor cortex (<45%) (Fig. 8A, blue). In the probe variant, decoding of the instructed action remained robust across all regions (MCM: 97.9%, MCL: 94.2%, SPL: 97.5%). Notably, the video action could now also be decoded from SPL with above-chance accuracy (52.9% at 2.3 s), whereas decoding from motor cortex remained at chance levels (Fig. 8A, gray). This contrasts with the no-probe variant, where the same visual input was present during intention but yielded no decodable information in any area, including SPL.

**Figure 8:**
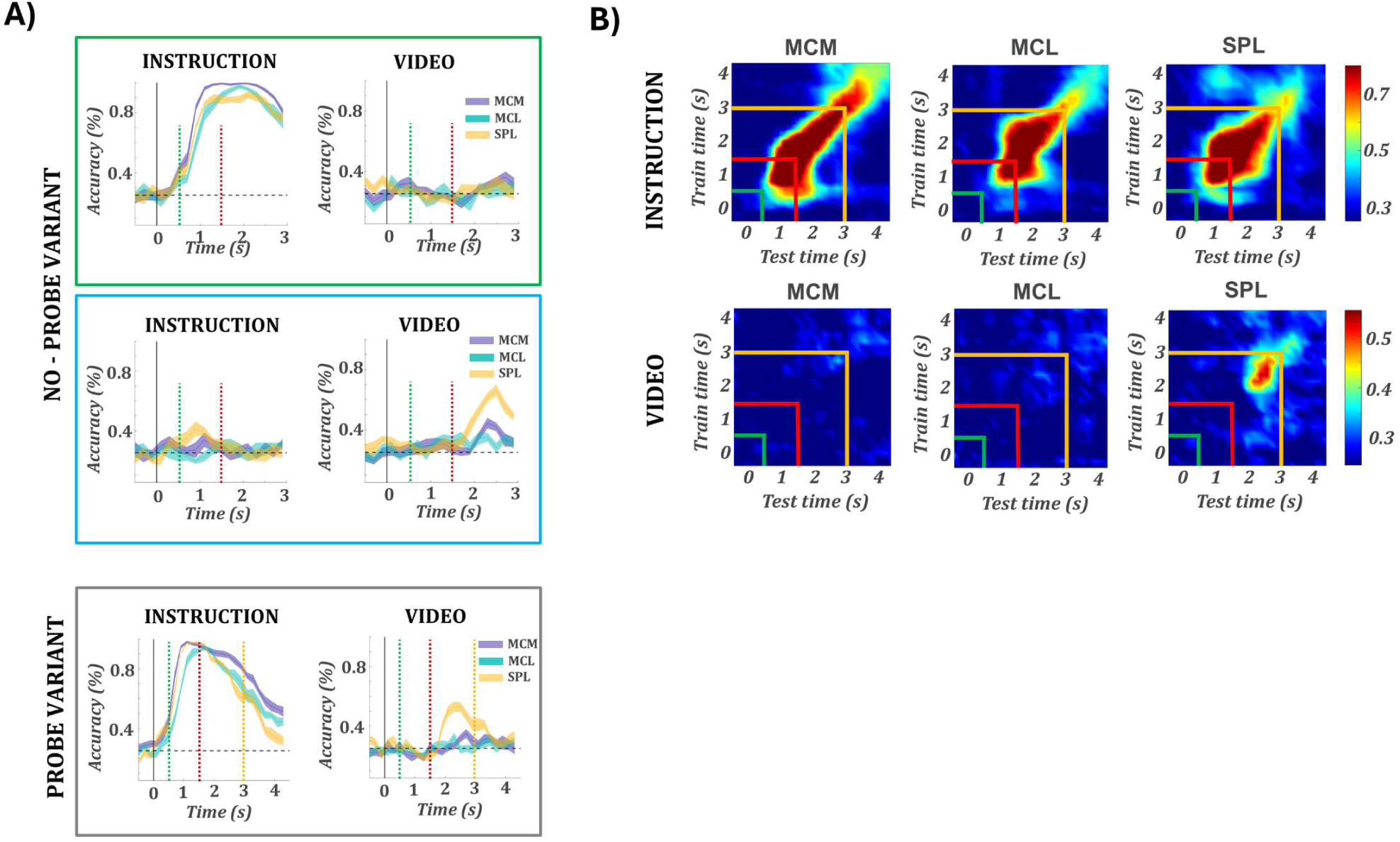
Dissociation tasks: decoding of instructed and observed actions. **(A)** Decoding of instructed (left) and video (right) actions. Green outline: no probe variant – instruction block; blue outline: no probe variant – observation block; gray outline: probe variant. For all decoding plots, colored lines show accuracy over time for MCM (purple), MCL (teal), and SPL (orange); shaded regions indicate ±SEM across cross-validation folds, and the dashed horizontal line marks chance level. In all plots, vertical lines indicate the hand cue (black), action cue (green), video onset (red), and probe onset (orange). (B) Heatmaps of LDA decoding accuracy across all train × test time windows for a given brain area (MCM, MCL, SPL). Top row: decoders trained on the instructed action; bottom row: decoders trained on the video action. Color indicates mean decoding accuracy across folds. The diagonal represents within-time decoding, whereas off-diagonal values reflect temporal generalization.

To rule out the possibility that decoding results were influenced by visual–motor congruency, we repeated all analyses in both the no-probe (intention and observation) and probe tasks using only incongruent trials. Decoding performance was virtually identical to that obtained when including all trials (Fig. S12A), indicating that population activity reflected genuine task-related representations rather than shared visual overlap. We also tested whether any units explicitly encoded the congruency between instructed and observed actions. Neither tuning nor decoding analyses revealed significant selectivity for conflict type across arrays, and decoding accuracy remained at chance in both task variants (Fig. S12B).

We next examined the temporal stability of the instructed and video action representations within the probe task using cross-time decoding (Fig. 8B). For the instructed action, all regions showed a strong and sustained diagonal, indicating stable decoding over time. The lack of broad off-diagonal generalization suggests that these representations were maintained but gradually reconfigured rather than held in a fixed subspace. SPL exhibited a slightly broader diagonal confined to the Go epoch, indicating that representations of the executed action were more temporally sustained during movement intention. In contrast, decoding of the video action was temporally restricted. Above-chance decoding emerged only in SPL and only within a short time window during the video presentation, indicating that SPL’s visual representations were brief and time-locked to sensory input, while motor cortex showed no reliable decoding.

To further characterize the structure of task representations, we trained classifiers to decode all 16 trial types from the dissociation task (2 actions × 2 hands × 4 conflict types; Fig. S13). In the no-probe intention block (Fig. S13A), decoding in MCM, MCL, and SPL revealed distinct clusters corresponding to the instructed action, consistent with selective encoding of executed movements. In the observation block (Fig. S13B), decoding accuracy dropped across all areas, and confusion matrices lacked systematic structure. In the probe task (Fig. S13C), motor cortex again showed clustered decoding aligned with the instructed action. In contrast, SPL exhibited a diagonal structure in the confusion matrix, indicating that it captured the full trial identity across all 16 conditions (peak accuracy: 49.6%), including both instructed and video features. This parallel representation emerged despite the fact that the task structure was nearly identical to the intention block of the no-probe version, differing only by the presence of a post-trial probe.

Together, these results reveal a gating mechanism in SPL. While motor cortex consistently reflected the instructed movement, SPL encoded observed (video) actions only when they were behaviorally relevant. This selective engagement occurred despite identical visual input across task variants, indicating that SPL’s visual selectivity was context-dependent rather than stimulus-driven. The absence of congruency or conflict effects confirms that decoding reflected genuine task-specific representations. In contrast, motor cortex maintained stable motor codes throughout. Overall, SPL dynamically allocated its representational resources according to behavioral relevance, supporting transient, task-dependent encoding of visual actions alongside stable motor representations.

## Discussion

We conducted the first systematic comparison of intended and observed actions using single-unit recordings in the human motor and posterior parietal cortex. In motor cortex, intended actions were robustly encoded, but passive observation failed to elicit mirror-like responses at the single-unit level. Still, population analyses revealed a weak but structured geometry during observation, partially aligned with the execution format and centered on the most strongly tuned features. In the superior parietal lobule, observed actions were reliably encoded, with most selective units maintaining shared tuning for action type across formats. These representations generalized across the population and were sensitive to behavioral relevance: when observed actions were not task-relevant, their encoding was suppressed. Our findings suggest that action observation engages distinct encoding schemes across cortical regions, reflecting a flexible, context-dependent system rather than a fixed mirror mechanism.

In motor cortex, the absence of mirror-like tuning at the single-unit level during passive observation contrasts with prior reports in nonhuman primates, where neurons in M1 and premotor areas exhibit consistent observation-driven responses^25–27^. However, population-level analyses revealed a more nuanced structure: neural activity projected into low-dimensional space revealed partially aligned trajectories between execution and observation, successful cross-decoding from observation to execution, and localized overlap in UMAP space, specifically for intention-tuned features (Figures 3-5). These results suggest that while single units do not overtly mirror observed actions, MC maintains a latent representational geometry during observation that reflects core aspects of the execution structure.

Jiang et al. (2020) ^28^ reported that executed and observed movements in monkey M1 and PMd occupy a shared subspace, with closely aligned population trajectories. In our recordings from human motor cortex, population overlap was more limited, confined to specific subregions and features strongly represented during intention. We also did not observe strong single-unit responses during passive observation, in contrast to reports in monkeys^25^, suggesting that human motor cortex may contribute less directly to action observation. Consistent with this, Rastogi et al. (2023)^29^ found that activity in human motor cortex was primarily structured by volitional state, with observation responses substantially weaker than those during attempted movement. Differences between species may partly reflect that our participants could not reproduce observed actions with matching kinematics, which could reduce overlap, but across human data the absence of robust single-unit mirroring appears to be a general finding. Our results extend this view by showing that, even without overt motor output, observation engages a small but structured latent representation in motor cortex, aligned with features strongly encoded during intention.

Our population-level findings are further supported by high-gamma LFP signals, which revealed robust and condition-congruent responses in both areas during observation fulfilling the classical criteria for mirroring (Fig. 6) and closely paralleling human fMRI results, that consistently report motor cortical activation during passive action viewing ^30^.The convergence across methods points to a representational gradient: while single units in motor cortex showed minimal tuning during observation, the population exhibited a weak but structured geometry that became more apparent at broader spatial scales. LFP, and by extension fMRI, may reflect subthreshold or spatially distributed synaptic activity. This interpretation is supported by evidence that the BOLD contrast mechanism correlates most strongly with LFPs (more so than with multi-unit activity) and primarily reflects synaptic input and intracortical processing rather than spiking output ^31^. Our results highlight a critical distinction: population-level metrics can reveal shared representational frameworks between formats even when overt mirror responses are absent at the single-neuron level.

In contrast to motor cortex, SPL exhibited robust encoding of observed actions, with a large proportion of selective units showing shared tuning to the same action across execution and observation (Fig 2E). The cross-format encoding was strongest for action type pointing to a higher-order representation of action identity. Such structure was also evident across the population. Neural trajectories were aligned across formats for matching actions; cross-decoding from execution to observation and vice versa was successful for action type; and UMAP projections revealed spatial overlap (Figures 4-6). These results suggest that SPL encodes a conceptual representation of action, generalizable across sensory format and resilient to contextual mismatch (e.g., when participants do not precisely reproduce the observed movement) not tied to specific motor output. Our findings converge with a growing body of work suggesting that PPC supports abstract action encoding. Aflalo et al. (2020)^24^ demonstrated shared population codes in human PPC for action verbs and visual action stimuli. Lanzillotto et al. (2020)^32^ showed that AIP neurons do not encode actions invariantly, but population activity allows reliable decoding across viewpoints and formats. Similarly, Chivukula et al. (2025)^22^ showed that somatosensory representations in PPC generalize across both experienced and observed touch. Across these studies, as in ours, PPC appears to encode not the physical parameters of an event, but the concept of the action itself. Our results extend this literature by showing that such conceptual generalization is supported by both single-unit tuning and aligned population geometry.

The dissociation paradigms allowed us to determine whether these abstract action representations in SPL are fixed or dynamically modulated. In the no-probe variant, SPL encoded only the instructed action, with no detectable representation of the concurrent videostream (Fig 7C, 7E). This dissociation confirms that the motor responses observed in our main task were not visually driven, ruling out the possibility that SPL activity merely reflects responses to visual input. In the probe variant (identical in sensory input but requiring participants to recall the video action on every trial) SPL encoded both the instructed and video actions (Fig 7D, 7F), whereas MC encoded only the instructed action in both task variants. To our knowledge, this is the first single-neuron evidence in humans of gating between motor and visual encoding in SPL. We show that observed features are not represented by default; their encoding emerges only when they are task-relevant. When motor and visual goals conflict, internally generated actions suppress irrelevant sensory input. The dissociation between motor output and visual input demonstrates that observed actions are not mirrored automatically but are flexibly gated by task demands.

Such task-dependent gating aligns with prior findings in nonhuman primates and human fMRI studies showing that sensory responses in PPC are modulated by cognitive context and behavioral relevance ^33^ and extends them by demonstrating flexible population-level reconfiguration at the level of individual neurons in human SPL. The role of SPL in this process is consistent with its proposed function within the frontoparietal multiple-demand (MD) network, a domain-general system implicated in cognitive control, goal-directed behavior, and adaptive task management ^34,35^. Rather than serving as a static relay of sensorimotor signals, PPC is increasingly viewed as a dynamic integrator whose representational structure is shaped by task goals. Models of attentional control further support this view, proposing that the dorsal frontoparietal network (including the intraparietal sulcus and superior parietal regions) allocates top-down attention and formulates predictions about incoming stimuli, while the ventral attention network centered on the temporoparietal junction and ventral frontal cortex) detects salient or unexpected events and redirects attention accordingly ^36^. These dynamics are closely related to the principle of mixed selectivity, whereby neurons encode combinations of task-relevant features across domains, a mechanism proposed to enable flexible and high-capacity representation in cognitive systems ^13–15^.

Our findings suggest that action understanding does not rely on automatic mirroring, but on a hierarchical organization in which motor cortex encodes intention-dominant signals and posterior parietal cortex flexibly integrates observed input with internal goals. This geometry-based dissociation points to context-sensitive transformations rather than reflexive resonance as the basis for linking perception to action. These conclusions are drawn from small cortical regions in two individuals with tetraplegia, raising the broader question of whether intact motor systems or other frontoparietal areas would reveal similar balances between intention and observation. Addressing such questions will be essential for testing whether the gating and geometric principles identified here reflect local properties or a general cortical strategy. In either case, our results highlight a framework in which human action representations emerge as flexible, context-dependent geometries embedded within the broader architecture of cognitive control.

## Methods

### Participants

Participants JJ and RD are right-handed males (ages 55 and 40) enrolled in a brain–machine interface (BMI) clinical trial (clinicaltrials.gov identifier: NCT01958086), approved by the institutional review boards of Caltech, Casa Colina Hospital, and UCLA. Participant JJ sustained a C4–C5 level spinal cord injury that occurred approximately 10 years prior to enrollment. He retains voluntary control of the eyes, head, and shoulders. Participant RD sustained a C3–C4 level spinal cord injury approximately 3 years prior to enrollment. He retains similar control of the eyes, head, and shoulders, and shows weak residual movements of the wrists and thumbs. Both participants were clinically stable at the time of participation. Presurgical functional MRI confirmed task-related activation in regions near the planned implant sites.

## Neural Recordings

Participant JJ was implanted in 2018 with two 96-channel Utah arrays targeting the left precentral gyrus (denoted JJ-MC) and superior parietal lobule (SPL) (denoted JJ-SPL). Participant RD was implanted in 2023 with four 64-channel Utah arrays targeting the left precentral gyrus (denoted RD-MCM, RD-MCL), SPL (denoted RD-SPL), and supramarginal gyrus (SMG). The SMG array in RD did not exhibit reliable task-related activity across sessions and was therefore excluded from all analyses. All arrays were 4 × 4 mm with 400 µm interelectrode spacing (Blackrock Microsystems). Neural signals were recorded using a 128-channel neural signal processor (NeuroPort System, Blackrock Neurotech). Multiunit activity (MUA) was sampled at 30 kHz and high-pass filtered at 750 Hz. Action potentials were detected using a threshold of –3.5 times the root mean square (RMS) of the high-pass filtered (250 Hz full bandwidth signal). Local field potentials (LFPs) were recorded continuously at a sampling rate of 1 kHz.

### Experimental setup

Experiments were conducted at Casa Colina Hospital and Centers for Healthcare for Participant RD, and at home for Participant JJ. In all sessions, participants remained seated in their motorized wheelchairs with their hands resting prone on a flat surface, in a well-lit room. A 30-inch LCD monitor was positioned directly in front of them. Stimulus presentation was controlled using the Psychophysics Toolbox^38^ for MATLAB (MathWorks).

## Action Intention and Observation Task

The experimental design consisted of a primary task performed under two cognitive conditions: intention and observation. In both conditions, participants viewed identical visual stimuli—a virtual hand performing actions—on a monitor positioned directly in front of them. Stimulus timing and content were matched across conditions. Eye position was continuously monitored using a Tobii eye tracker to confirm that participants attended to the stimuli, although no fixation requirement was imposed.

In the intention block, participants were instructed to internally generate the cued action using the specified hand while looking at the screen. Participant JJ remained physically still but volitionally intended the instructed action. Participant RD, who retained partial motor function, was instructed to overtly perform the cued action. Although his movements did not always replicate the precise kinematics of the video stimuli due to physical limitations, we allowed naturalistic execution and instructed him to maintain consistency in the type of response across trials and sessions. In the observation block, participants passively viewed the same videos while remaining still. They were instructed to observe the actions without intending any movement.

Each trial began with a 0.5-second inter-trial interval (ITI), followed by a 0.5-second hand cue. The hand cue consisted of a static image of a virtual left or right hand holding a small parallelogram object between the thumb and index finger, indicating the instructed effector. This was followed by a 1-second symbolic action cue, presented as an overlaid arrow indicating both the action type and direction. Arrow shape specified the action: straight for sliding, curved for lifting, and circular for rotating. The endpoint of the arrow indicated the direction of movement (leftward or rightward). For sliding and lifting, this corresponded to a horizontal displacement of the object to the left or right, while for rotation it corresponded to a clockwise or counterclockwise turn. For consistency, we refer to this factor abstractly as left/right direction throughout the manuscript. The disappearance of the symbol served as the go cue. Immediately afterward, the static frame of the hand transitioned into a 1-second video showing the hand performing the cued action in the indicated direction. The task followed a fully crossed 2 (hand) × 3 (action) × 2 (direction) design, resulting in 12 unique conditions. Each participant completed 12 repetitions per condition, per block, in each session. Participant JJ completed five sessions, yielding a total of 421 units in MC and 326 units in SPL, aggregated across sessions. Participant RD completed six sessions, yielding 441 units in MCM, 479 units in MCL, and 532 units in SPL.

## Dissociation Task Variants

To examine how task demands and cue congruency shape action representations, Participant RD completed two additional experiments with an observation phase that is dissociated from the instruction phase. These experiments were performed only with RD, who retained partial motor function and was able to overtly execute the instructed actions. This was essential to ensure correct task performance in the dissociation paradigm, where precise execution of the cued action—despite incongruent visual input—was required. To reduce complexity, we fixed the direction of movement to rightward and used only two action types: *slide* and *rotate*, performed with either the left or right hand. This yielded a 2 (action) × 2 (hand) design. For rotation, the rightward direction corresponded to a clockwise turn, while for sliding it corresponded to a rightward displacement.

Each trial began with a 0.5-second inter-trial interval (ITI) displaying a fixation cross. This was followed by:

- a 0.5-second hand instruction screen, indicating the instructed effector (left or right hand),
- a 1-second symbolic cue, showing an overlaid arrow that specified the instructed action type (slide or rotate),
- and a 1-second go phase, during which a video of a hand performing an action was displayed.

The action in the video could be congruent or incongruent with the instructed action. Specifically, the trial could present one of four cue–video pairings:

1. Fully congruent (same action and hand),
2. Action incongruent (same hand, different action),
3. Hand incongruent (same action, different hand), or
4. Action and hand incongruent (different action and hand).

The participant was instructed to perform the cued action with the specified hand concurrently with the video playback, regardless of what was shown.

## Dissociation–No-Probe Task

This version included two blocks: an intention block and an observation block. In the intention block, the participant executed the instructed action while watching the video. In the observation block, the participant passively viewed the same videos, with no movement or intention. No response was required after the trial. This allowed us to dissociate encoding of instructed versus observed actions.

## Dissociation–Probe Task

This version was identical in structure but added a 1-second probe screen immediately after the video. On each trial, the probe queried either the hand or the action shown in the video. Two icons were presented (left vs. right hand, or slide vs. rotate), and the participant was required to make a saccade to the correct icon to report what they had observed. This response was recorded using the eye-tracking system. By forcing explicit recall of the observed action or effector, while the participant simultaneously performed the instructed action, this task ensured that both streams of information were behaviorally relevant.

### Spatial Distribution of Gaze

To visualize spatial gaze patterns during the experiment, we computed two-dimensional heatmaps of gaze position separately for the intention and observation blocks, combining data across all runs and sessions. Gaze coordinates from all trials were concatenated, and two-dimensional histograms were computed using a fixed grid (100 × 100 bins) spanning the full range of observed horizontal and vertical gaze values. The resulting gaze density distributions were smoothed using interpolation for visualization, yielding continuous heatmaps that reflect the spatial concentration of gaze over the course of the experiment.

## Data Preprocessing

### Single Unit Activity (SUA) and Multiunit Activity (MUA)

Each detected waveform consisted of 48 samples (1.6 ms total), including 10 samples before threshold crossing and 38 samples after. Single-unit and multiunit activity were sorted using Gaussian mixture modeling applied to the first three principal components of the waveform shapes^14^. In addition to spike assignments, the sorting procedure provided a quality factor ranging from 1 to 4, determined by (1) the percentage of interspike intervals shorter than 3 ms, (2) the signal-to-noise ratio of the mean waveform, (3) the projection distance between clusters, (4) the modified coefficient of variation of the interspike intervals (CV2), and (5) the isolation distance of each cluster. Units with a quality factor of 1 or 2 were considered well-isolated, whereas those with a factor of 3 or 4 were classified as multiunit activity. ^14^ After spike sorting, net average responses were computed in 100 ms bins by subtracting baseline activity (−500 to 0 ms before stimulus onset) from the post-stimulus activity (0 to 3000 ms after stimulus onset) on a trial-by-trial basis.

### Local Field Potential (LFP)

Line noise was suppressed using a combined spectral and spatial filtering approach that preserves underlying neural signals while attenuating non-neural components^39^. The data were then high-pass filtered above 2 Hz using a zero-phase infinite impulse response (IIR) filter. Trials in which the broadband signal amplitude exceeded two standard deviations from the session mean were excluded from further analysis. High-gamma (60–120 Hz) power was estimated using Morlet wavelet convolution with a 7-cycle resolution^40^. To minimize edge artifacts introduced by filtering and wavelet convolution, the first and last 100 ms of each trial were discarded. Power was normalized within each trial by dividing the time–frequency power at each frequency by the mean power at that frequency during the 500 ms pre-stimulus baseline.

## Quantification and Statistical Analysis

### Statistical Comparison to Baseline

To identify task-modulated activity, we compared post-stimulus responses to baseline on a per-condition basis. For high-gamma analyses, this was performed at the channel level, for spiking activity, at the unit level. Neural responses (spike rates or high-gamma power) were binned into overlapping 200 ms windows with a 100 ms step size, starting at stimulus onset. For each bin, a paired t-test compared the binned activity to the mean activity during the 500 ms pre-stimulus baseline across trials. Statistical significance was assessed using a Bonferroni-corrected threshold (α = 0.05 / number of bins). A channel or unit was classified as significantly responsive to a condition if it exhibited at least three consecutive time bins with significant deviation from baseline. This analysis was performed separately for each brain region, condition, and signal type.

### Latency

For each condition and each significantly responsive unit, response latency was defined as the center of the first time bin within the earliest sequence of three consecutive bins showing significant modulation relative to baseline. This latency reflects the earliest consistent deviation from baseline activity. To assess regional differences in response timing, we compared latency distributions across brain areas using one-way ANOVAs, conducted separately for each participant and for each task format (intention and observation). Each unit contributed a single latency value per condition, and group-level comparisons tested for significant differences in peak response timing between motor and parietal regions.

### Tuning Analysis

To quantify selectivity for task variables, we performed a time-resolved three-way ANOVA separately for spiking activity (multi– and single-unit) and high-gamma (HG) power. For each unit or channel and for each 500 ms time bin, we computed the mean response across time. A three-way ANOVA was then used to test for main effects of action type (rotate, slide, lift), effector (left vs. right hand), and movement direction (leftward vs. rightward), as well as their three-way interaction. For the dissociation tasks, in which movement direction was held constant, a two-way ANOVA was performed with action and hand as factors. Trial labels were parsed from condition names into categorical variables corresponding to each factor. Significance was assessed using a Bonferroni-corrected threshold (α = 0.05 / number of units or channels). A unit or channel was classified as selectively tuned to a main effect (e.g., action type) if the corresponding factor reached significance in the absence of a significant interaction. Tuning to a combination of factors was labeled as an interaction effect. This analysis was performed independently for each time bin and brain region, and the number of significantly tuned units or channels was tracked across time for each tuning category.

### Overlap of Task-Relevant Units Across Formats

To assess the distribution of task-related neural responses across formats, we identified task-responsive units independently for the intention and observation blocks. A unit was classified as task-responsive if it exhibited a significant increase in activity relative to baseline for at least one task condition (see *Statistical Comparison to Baseline*). This analysis was performed separately for each brain region and format. We then quantified the degree of overlap between formats by computing, for each region, the number of units responsive exclusively during intention, exclusively during observation, or during both. These distributions were used to evaluate the extent to which task-related neural activity was shared or format-specific across the two conditions.

### Linear Model Analysis

We classified units based on their selectivity to task variables across observation and intention conditions. We implemented a structured model comparison framework similar to that described by Chivukula et al., 2025^22^. For each neuron, firing rates were averaged over a fixed task window (1–2 s after trial onset) for each of the 12 observation and 12 intention conditions. These 24 condition-averaged responses were then combined into a single dataset for that unit, and a series of linear regression models was fit to predict neural responses based on different combinations of experimental factors (hand used, action type, movement direction) and format-specific terms (intention vs. observation).

(1) a null model including only a constant term (*Unselective*)
(2) a fully shared model including action, hand, and direction as predictors with the same weights across formats (*Invariant*)
(3–5) models with shared tuning to a specific task variable: action, hand, or direction (*Action*, *Hand*, or *Direction*)
(6) an additional model in which “action” was redefined as six distinct action × direction combinations (*Action (6)*);
(7) a fully shared model with all main effects and interactions (*Mixed*)
(8) a format-specific model with separate parameters for intention and observation (*Idiosyncratic*)
(9–10) models including task features only for intention or only for observation trials (*Single format*)

Model performance was assessed using five-fold stratified cross-validation based on condition labels, and each unit was assigned to the model with the highest cross-validated R² value. To determine whether the observed R² exceeded chance, we implemented a permutation test in which the neural responses were randomly shuffled across trials while keeping the design matrix fixed. For each unit, we computed a null distribution of R² values (1,000 permutations) and derived a one-tailed p-value based on the proportion of null R² values exceeding the true R². P-

values were corrected for multiple comparisons across units using the Benjamini–Hochberg false discovery rate (FDR) procedure (q = 0.05). Units were considered selective if they met all of the following criteria: (1) cross-validated R² > 0.01, (2) permutation-derived p-value < 0.05 after FDR correction, and (3) the best-fitting model was not the null model. Units that did not meet all three criteria were classified as unselective.

## Population Analysis

### Representational Similarity Analysis

To evaluate the structure of neural representations across intention and observation, we performed a cross-validated representational similarity analysis (RSA), separately for each task variable: action type (3 levels), effector/hand (2 levels), and movement direction (2 levels). Trial labels were regrouped accordingly, and only task-relevant units in both formats, defined as those that were neither unselective nor best fit by single-format models in the linear model-based tuning analysis (see *Linear Model Analysis*), were included. All analyses were conducted independently per brain region. RSA was computed both *within format* (intention–intention and observation–observation) and *across format* (intention–observation), using the same framework across all comparisons. Neural responses were extracted from a fixed 1–2 s post-stimulus window, and each trial was reshaped into a single feature vector by concatenating the time and unit dimensions. For each of 500 random splits, trials were divided into halves within each condition. Condition-averaged activity vectors were computed independently for intention and observation trials, and Pearson correlations were calculated between all pairs of vectors across formats, resulting in a condition-by-condition cross-format RSA matrix per split. These matrices were averaged to obtain a final similarity matrix. To test whether the observed structure reflected meaningful condition-specific information, we compared the similarity between matched and unmatched conditions by computing the mean diagonal and off-diagonal values of each RSA matrix. The difference between these values (diagonal – off-diagonal) served as a measure of structure strength. To generate null distributions, we repeated the same procedure after shuffiing condition labels independently within each format prior to trial splitting. Statistical significance was assessed by comparing the observed difference to the empirical null distribution from shuffied data, separately for the cross-format matrix and for each within-format matrix (intention and observation). In addition, we computed the full RSA matrices and permutation-matched nulls for each format and task variable. Mean diagonal values were extracted as a measure of within-condition reliability and representational consistency.

To assess the robustness of the RSA results, we conducted control analyses using four alternative time windows (0–1 s, 0.5–1 s, 1.5–2.5 s, and 2–3 s). For each window, we recomputed RSA matrices for intention, observation, and cross-format comparisons, and then quantified similarity to the main analysis (1–2 s window) by computing the Pearson correlation between the lower triangles (excluding the diagonal) of each matrix and the corresponding 1–2 s RSA matrix. This analysis provided a direct measure of how stable the representational structure was across different temporal windows.

We also repeated the RSA using all recorded units, rather than only task-relevant ones, and computed matrix correlations between the RSA results from all units and those from relevant units only for each comparison type (intention, observation, and cross-format).

### Linear Discriminant Analysis (LDA) Within Format

We assessed whether neural population activity encoded task-relevant information within each format, by training LDA classifiers using a time-resolved, cross-validated decoding framework. The analysis was performed separately for each brain area and each classification level. For the main task, classifiers were trained to distinguish among: (1) the 12 fully crossed task conditions (hand × action × direction), and (2) individual task variables: action type (3 levels), effector/hand (2 levels), and movement direction (2 levels). Trials were relabeled accordingly, and each classification problem was evaluated independently. For the dissociation tasks, decoding was performed in two complementary ways. First, classifiers were trained to distinguish among all 16 trial types, defined by the 2 (action) × 2 (hand) × 4 (conflict type) design. Second, to separately assess encoding of the instructed and observed actions, trials were relabeled based on the 2 (action) × 2 (hand) combinations corresponding to either the instructed cues or the observed video, and decoding was performed independently for each. For all analyses, neural responses were binned in 100 ms steps, and decoding was performed using non-overlapping 200 ms windows by averaging adjacent bins. Ten-fold cross-validation was used. Within each fold, principal component analysis (PCA) was applied to the training data across all time bins to reduce dimensionality. The number of components was selected to capture 95% of the variance, with an upper limit of 50 components. Both training and test data were projected into this reduced-dimensional space. LDA classifiers were trained on the reduced training data and evaluated on the held-out test set for each time window. Classification accuracy was computed per fold and averaged across folds to yield time-resolved performance curves for each brain region and classification level. For each region, the time window with the highest decoding accuracy was identified, and confusion matrices were computed at that time point. Confusion matrices were normalized by row and visualized as percent classification accuracy. To evaluate the consistency of decoding performance across sessions, we additionally ran the same decoding analysis separately within each session. This was done for each of the three individual task variables (action, hand, direction), and the maximum decoding accuracy per session was extracted. This allowed us to assess whether the results observed in the concatenated analysis were driven by any session-specific peaks or drops in performance.

### Cross – Time Decoding

To evaluate the temporal stability of neural representations, we performed cross-time decoding within the probe variant of the dissociation task, separately for each brain area and for classifiers trained on the instructed or video actions. The analysis followed the same preprocessing and cross-validation procedures described above. For each array, neural activity was binned in 100 ms steps, and non-overlapping 200 ms windows were created by averaging adjacent bins. Within each fold of a ten-fold cross-validation, principal component analysis (PCA) was computed on the training data across all time bins, and both training and test data were projected into the same reduced-dimensional space using the coefficients derived from the training set (95% cumulative variance threshold, maximum 50 components). LDA classifiers were then trained to discriminate either the instructed or the video actions at each time window and tested on all other time windows, generating a two-dimensional matrix of decoding accuracy (train × test time). This procedure was repeated for all folds, and decoding accuracies were averaged across folds to yield cross-temporal generalization matrices for each array. The diagonal of the matrix reflects standard within-time decoding, while off-diagonal values quantify the degree to which representations generalize across time. Sustained off-diagonal accuracy indicates temporally stable population codes, whereas narrow, diagonal patterns correspond to dynamic, time-specific encoding.

### LDA Within Format – LFP

To decode task variables from LFP, we extracted high-gamma (HG) power following the methodology described in Bouchard et al. 2013^41^. Raw LFP signals were re-referenced using common average referencing (CAR) and filtered using eight Gaussian-like bandpass filters, with logarithmically spaced center frequencies between 73 and 144 Hz. The bandwidth for each filter was scaled semi-logarithmically at 20% of the center frequency. The analytic amplitude for each filtered signal was computed using the Hilbert transform, and the resulting envelopes were downsampled to 100 Hz. Amplitude envelopes were then z-scored per channel across all timepoints and trials, and outlier suppression was applied using a hyperbolic tangent function. To reduce dimensionality across the eight frequency bands, we performed singular value decomposition (SVD) on the concatenated envelope matrix (channels × time × frequency) and retained the first singular vector as the unified HG estimate per channel. We identified task-relevant channels by comparing HG power during a task window (1–2 s post-stimulus) against a baseline window (0–0.5 s) using paired t-tests per channel and included only those with significant differences (p < 0.05). LDA decoding was then performed separately for each brain area and task variable (action type, effector/hand, movement direction), as well as for the full 12-condition design. Decoding was performed in 500 ms non-overlapping windows across the trial duration (−0.5 to 3 s), using 10-fold cross-validation. Classification accuracy was computed per time bin, and peak decoding performance was visualized alongside confusion matrices derived from the best-performing time window.

### Cross – Format Decoding

We evaluated whether neural population representations generalized across formats, by training LDA classifiers on trials from one format (intention or observation) and testing them on the other. This analysis was performed separately for each brain region and each task variable: action type (3 levels), effector/hand (2 levels), and movement direction (2 levels). Decoding was conducted in both directions: training on intention and testing on observation, and vice versa. Neural activity was binned in 100 ms steps, and decoding was performed per time bin. For each time bin in the training format, PCA was applied to the full training dataset (all time bins and trials), and the top 50 components were retained. Training and test data were projected into this common PCA space. An LDA classifier trained on each individual training time bin was then evaluated on all test time bins, yielding a full train × test time bin decoding accuracy matrix. This matrix captures temporal generalization and reveals whether neural representations aligned across different time points between formats. To assess statistical significance, we performed permutation testing by randomly shuffling test set labels across 1000 iterations and recomputing decoding accuracy. For each train–test pair, a p-value was computed as the proportion of permuted accuracies greater than or equal to the observed accuracy. Significance thresholds were corrected for multiple comparisons using Bonferroni adjustment.

### Cross – Format Decoding LFP

To evaluate whether HG activity encoded similar task-related information across formats, we performed cross-format decoding using the same HG signal described in the within-format LFP decoding analysis (see *LDA Within Format – LFP*). For each brain area and task variable (action type, effector, direction), a linear discriminant analysis (LDA) classifier was trained on single-trial HG activity from one format (e.g., intention) and tested on data from the other (e.g., observation), and vice versa. HG power was averaged within non-overlapping 500 ms windows spanning −0.5 to 3 s, yielding one feature vector per trial per time bin. Decoding accuracy was computed for all train × test time bin combinations, resulting in a time-resolved decoding matrix for each direction. Statistical significance was assessed by generating null distributions through 1000 permutations of the test labels. Empirical p-values were computed for each timepoint pair and corrected for multiple comparisons using a Bonferroni threshold.

### PCA Trajectory Analysis

To visualize the temporal evolution of neural population activity during each task, we performed PCA separately for each brain region and task variable (action type, effector/hand, or movement direction). Only units classified as task-relevant in both formats based on the linear model analysis (i.e., not unselective or single format) were included. Neural responses were extracted from a time window spanning –0.5 to 2.5 s relative to trial onset, using 100 ms binning. The PCA was applied jointly to intention and observation data. For each trial, neural activity was grouped by condition (e.g., action identity) and format (intention or observation). Trials were averaged within each condition and format, resulting in a set of condition-averaged response matrices of size time × units with the number of matrices equal to the number of task levels (e.g., 3 for action, 2 for hand) multiplied by 2 formats. These matrices were concatenated across time and condition, and the resulting (conditions × time) × units matrix was used as input to PCA. The top three principal components were retained, and each condition’s time series was projected into this low-dimensional space. The resulting trajectories were plotted separately for intention and observation to compare their temporal geometry. In addition, we quantified the variance explained by the top three components in three ways: based on the PCA decomposition of the combined dataset, and separately for intention and observation by computing the variance captured within each format after projection into the shared PCA space.

### Procrustes Analysis

To compare the geometry of neural trajectories between formats, we applied Procrustes analysis to the condition-averaged neural trajectories in PCA space. This analysis was performed separately for each brain region and task variable (action type, effector/hand, or movement direction). For each condition within a task variable (e.g., “lift” or “right hand”), we extracted one trajectory from the intention format and one from the observation format, both represented in the space defined by the top three principal components computed from the combined dataset. Procrustes alignment was then used to align the 3D trajectories of the two formats. This transformation computes the optimal translation, rotation, and isotropic scaling that minimizes the Frobenius norm between the two trajectories. A Procrustes distance was computed for each condition, providing a measure of geometric dissimilarity between formats. Lower values indicate greater similarity in the temporal structure of neural activity between intention and observation for that condition. As a control, we repeated the PCA and alignment procedure using all recorded units, and computed Procrustes distances for all brain regions and task variables.

### UMAP Visualization of Neural Representations

To visualize the structure of trial-level neural activity in a low-dimensional space, we applied Uniform Manifold Approximation and Projection (UMAP) separately for each brain region. Only task-relevant units (identified via linear model analysis) were included. Neural responses were averaged over a 1–2 s window following stimulus onset. Trials from the intention and observation formats were concatenated and projected jointly. UMAP was performed using the cosine distance metric with 15 nearest neighbors and a minimum distance of 0.1. The resulting two-dimensional embeddings were used to visualize the structure of population activity across task conditions and formats. To visualize the spatial distribution of each condition in the UMAP space, we computed a separate 2D ellipse for each condition and format combination. The center of each ellipse was defined as the empirical centroid of the UMAP coordinates for that group (i.e., the mean across all trials). The shape and orientation of the ellipse were derived from the empirical 2D covariance matrix of the points. We used the eigenvectors and eigenvalues of this matrix to construct an ellipse representing the major and minor axes of the group’s spread. This approach provides an interpretable summary of the distribution of trials in the reduced space, allowing visual assessment of separation or alignment between formats.

## Data Availability

All data produced in the present study are available upon reasonable request to the authors

## Acknowledgements

We would like to express our gratitude to participants N1 and N2 for making this research possible. This work was supported by National Eye Institute grant UG1EY032039, Tianqiao and Chrissy Chen Brain-Machine Interface Center at Caltech, Swartz Foundation, and Boswell Foundation. VB is supported by Fonds Wetenschappelijk Onderzoek (FWO) grant 1264926N.

## Contributions

VB, JG, and RAA conceived and designed the experiment. VB performed the recordings, data analysis and wrote the manuscript. JG and RAA supervised and guided the study. ERR, AAB, and CL provided clinical support. VB, JG, RAA, KP acquired funding. KP and RAA were responsible for project administration. All authors reviewed and edited the manuscript.

