## Supplementary Material for "Hierarchical and Context-Dependent Encoding of Actions in Human Posterior Parietal and Motor Cortex"

**Supplementary Figures:**

**Figure S1:** **Gaze heatmaps during intention and observation.**

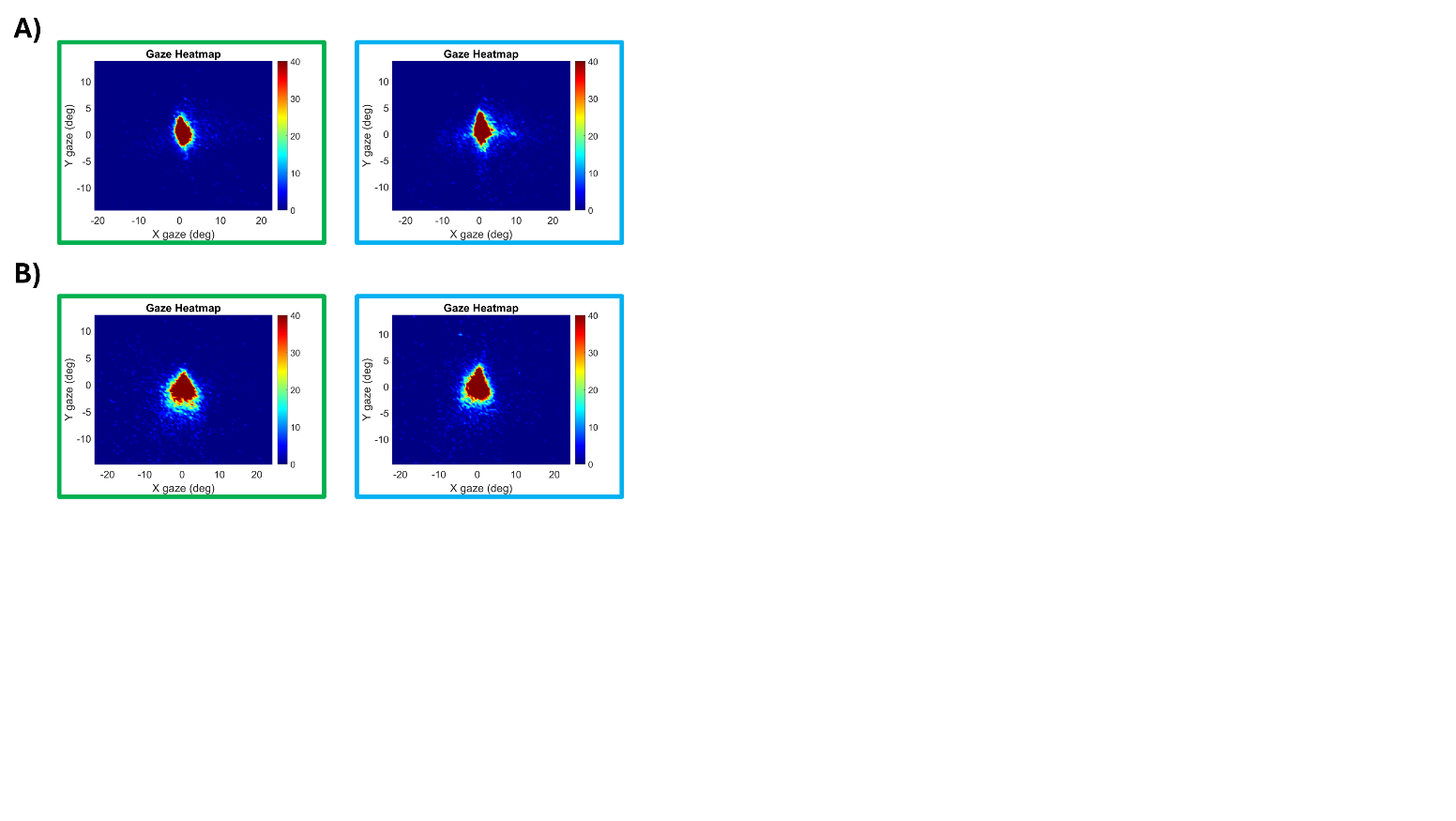

**(A**) Participant JJ**. (B)** Participant RD. Heatmaps show the 2D distribution of gaze positions (in degrees of visual angle) across all trials. Left panels (green outline) correspond to the intention format; right panels (blue outline) to the observation format. Across both participants and formats, gaze remains consistently centered, confirming fixation compliance throughout the experiment. Color scale indicates gaze density.

**Figure S2: Linear model analysis including the expanded six-action model.**

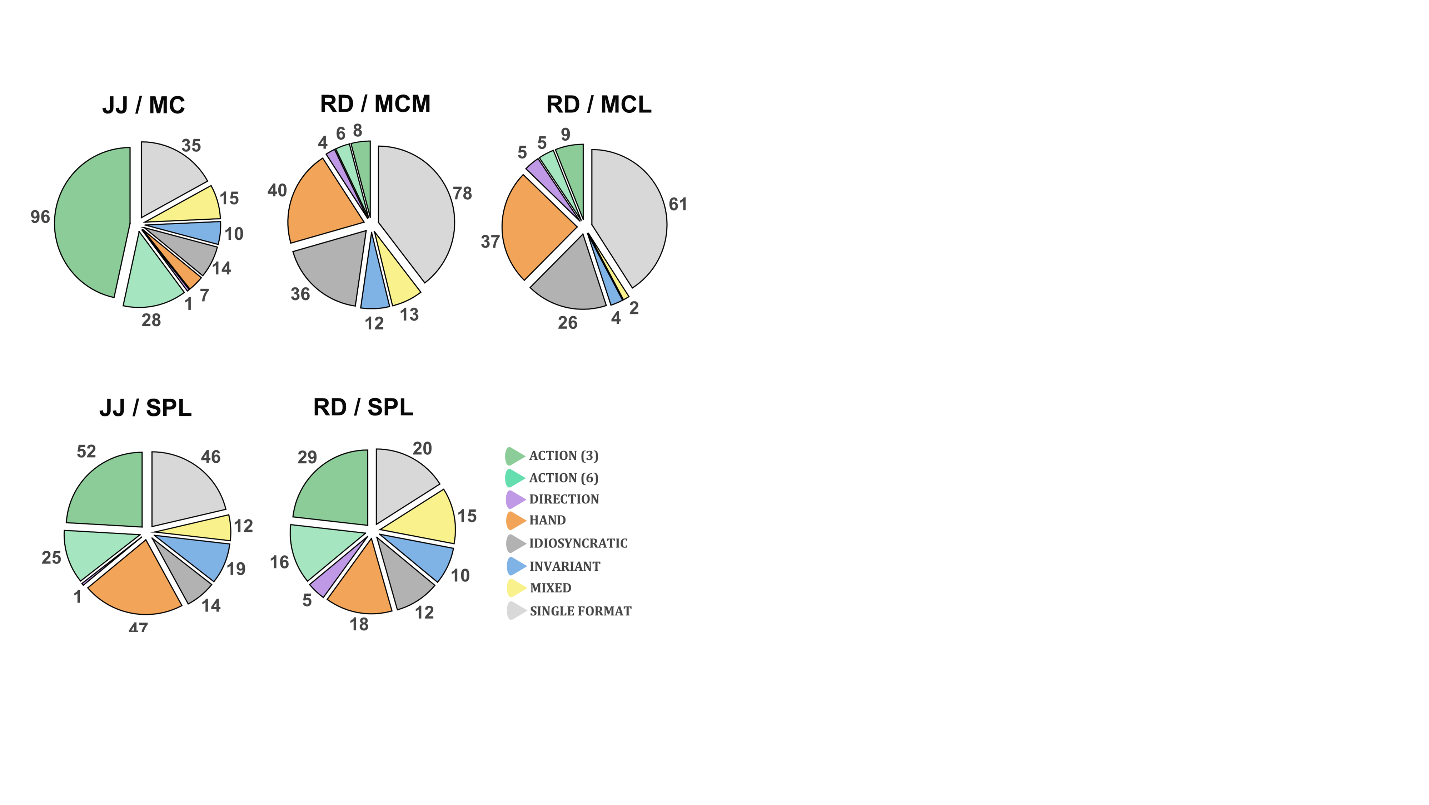

Distribution of best-fitting linear models across brain areas. Units are assigned to their best-fitting model based on cross-validated R². Categories include shared action (3) (dark green), shared action (6) (light green), shared hand (orange), shared direction (purple), invariant (blue), mixed (yellow), idiosyncratic (dark gray), and single-format (light gray). Unselective units are not shown.

**Figure S3:** **Within-format representational similarity analysis (RSA)**

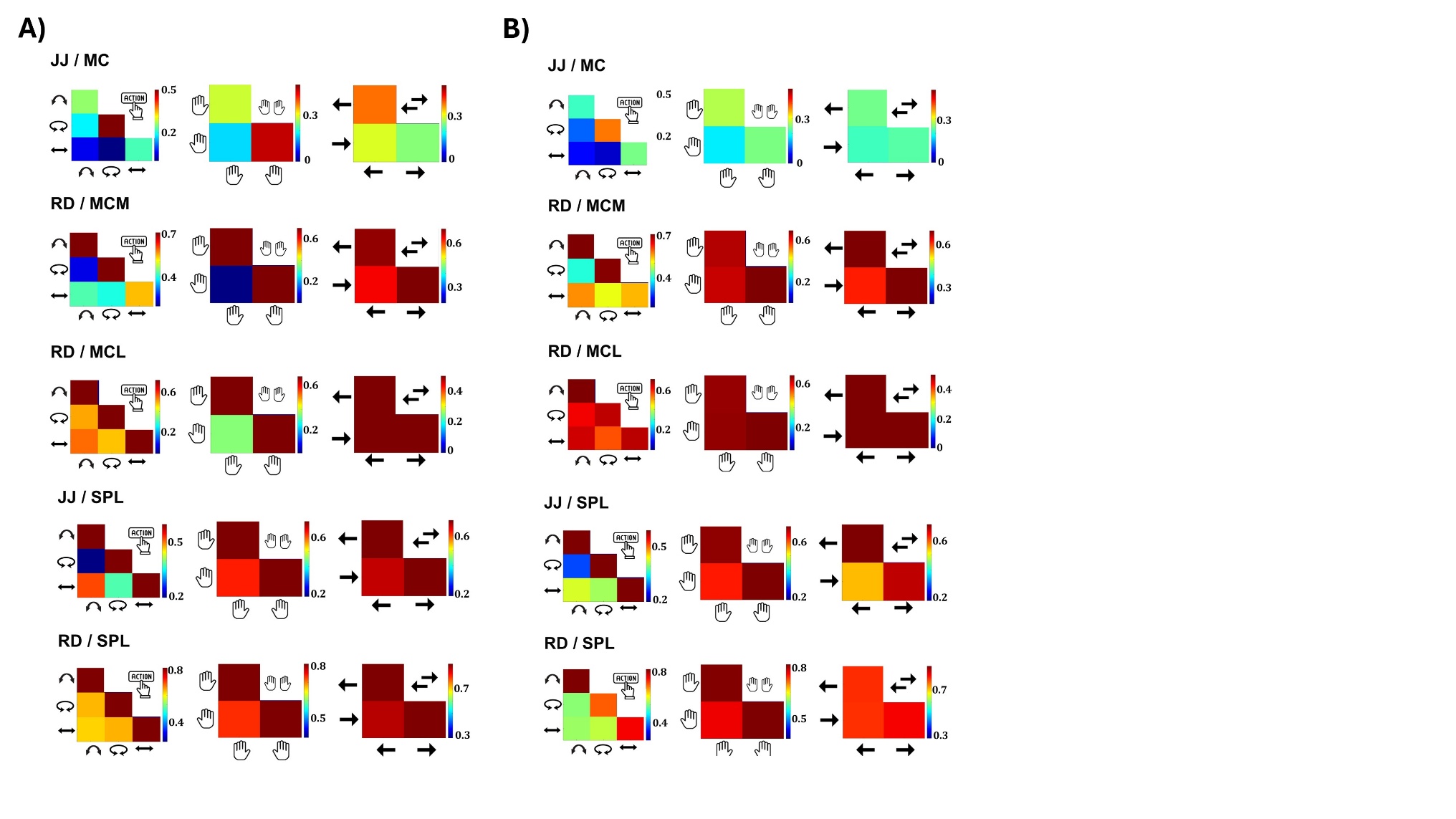

**(A)** RSA heatmaps computed within intention. Columns correspond to action (left), hand (middle), and direction (right). Each matrix shows the pairwise correlation structure between condition-specific population activity vectors. Similar patterns to the cross-format RSA emerge: JJ/MC shows high correlations for the rotation action, and RD/MCM shows structured representation for right hand, rotation, and lift actions. Additional structure appears for left hand in MCM and right hand in MC/RD. MCL now exhibits clear diagonals for both hand and action. SPL arrays in both JJ and RD show structured representations for action. **(B)** RSA heatmaps within observation. Action selectivity patterns remain visible in JJ/MC and RD/MCM. However, separability by hand is lost in all arrays. MCL shows globally high correlations across all features, indicating poor condition discrimination. Only SPL arrays show well-structured diagonals for action types during observation.

**Figure S4:** **Control analysis for RSA matrix stability and unit selection.**

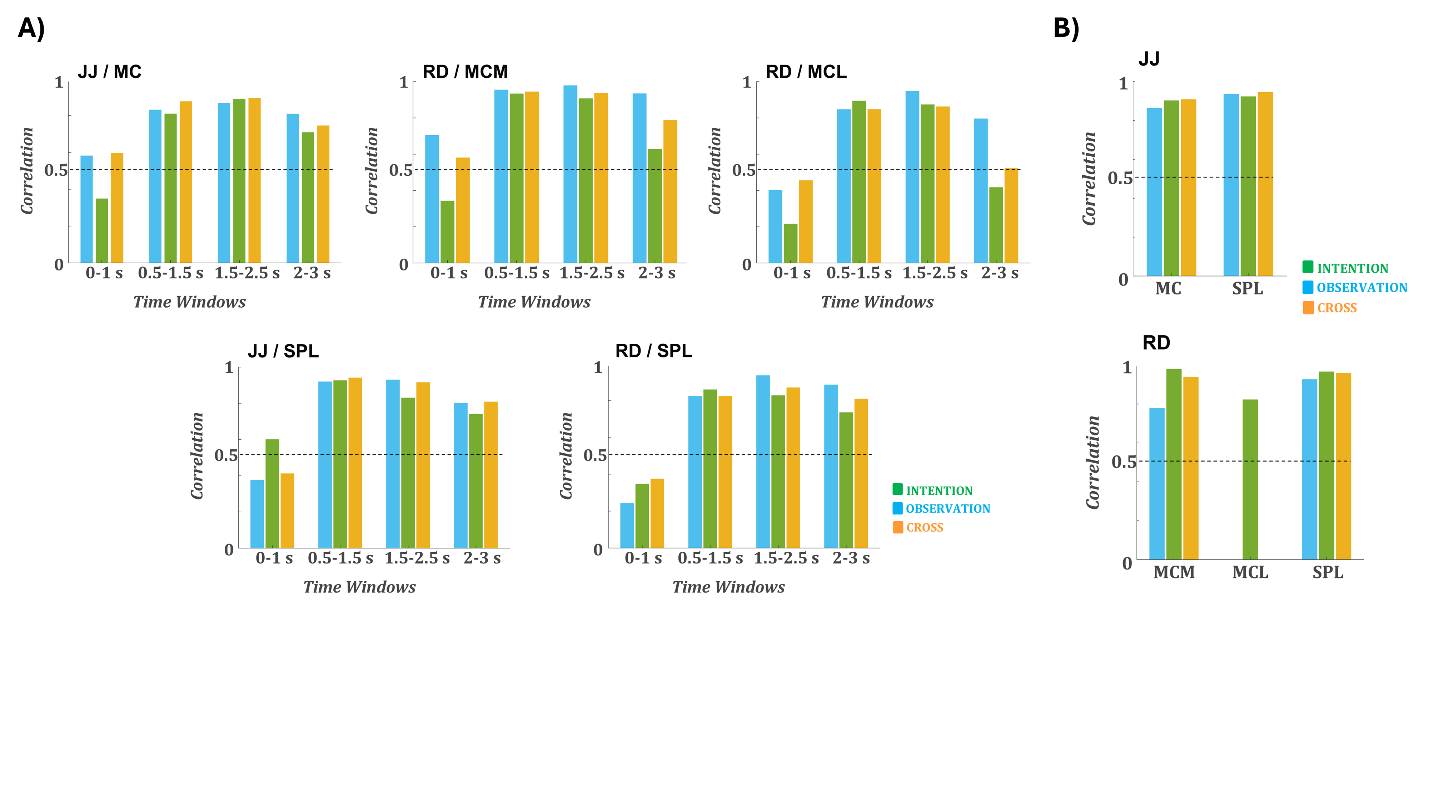

**(A)** Robustness of representational structure across time windows. Bars show correlations between the lower triangle of the 12-condition RSA matrices computed in the main analysis window (1–2 s) and alternative windows: 0–1 s, 0.5–1.5 s, 1.5–2.5 s, and 2–3 s. Colors indicate format: within observation (blue), within intention (green), and cross-format (orange). Correlation remains high for all windows except the earliest (0–1 s), confirming the stability of the representational structure over time. **(B)** Effect of unit selection. Bars show the correlation between RSA matrices computed using all recorded units and those computed using only task-relevant units (used in the main analyses), for each format. Correlations are high across all arrays and formats, confirming that the main results are not driven by unit selection. MCL, the correlation is lower, but this reflects the fact that both RSA matrices (all and task-relevant) lack meaningful structure, resulting in low but non-contradictory values.

**Figure S5:** **Decoding of all task conditions within format.**

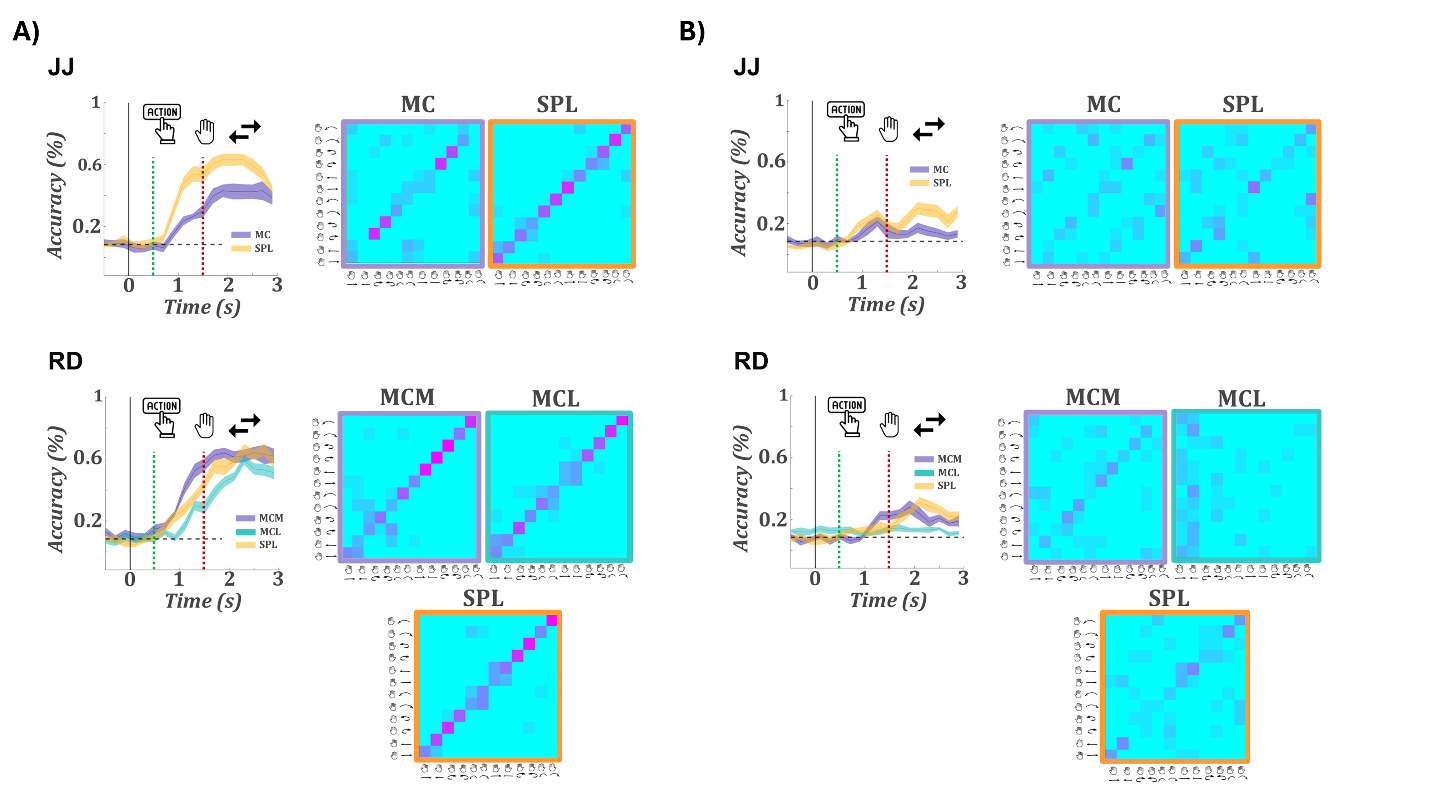

**A)** Intention. Left: time-resolved LDA decoding accuracy for all 12 task conditions (2 hands × 3 actions × 2 directions). Top: JJ (MC, SPL); bottom: RD (MCM, MCL, SPL). Curves show average accuracy across cross-validation folds; shaded regions indicate ±SEM. Green and red dashed lines mark the symbolic action cue and go cue, respectively. All arrays achieve high classification accuracy, with SPL consistently performing best. Right: corresponding confusion matrices show that most misclassifications occur between different directions of the same action, indicating that action identity and hand are more robustly represented than direction. **B)** Observation. Same format as A. Decoding accuracy is substantially lower across all arrays. SPL outperforms other regions in both participants, but peak accuracy remains below 40%. Confusion matrices show reduced separability between conditions, consistent with weaker observation-related encoding.

**Figure S6:** **Session-wise decoding accuracy within format.**

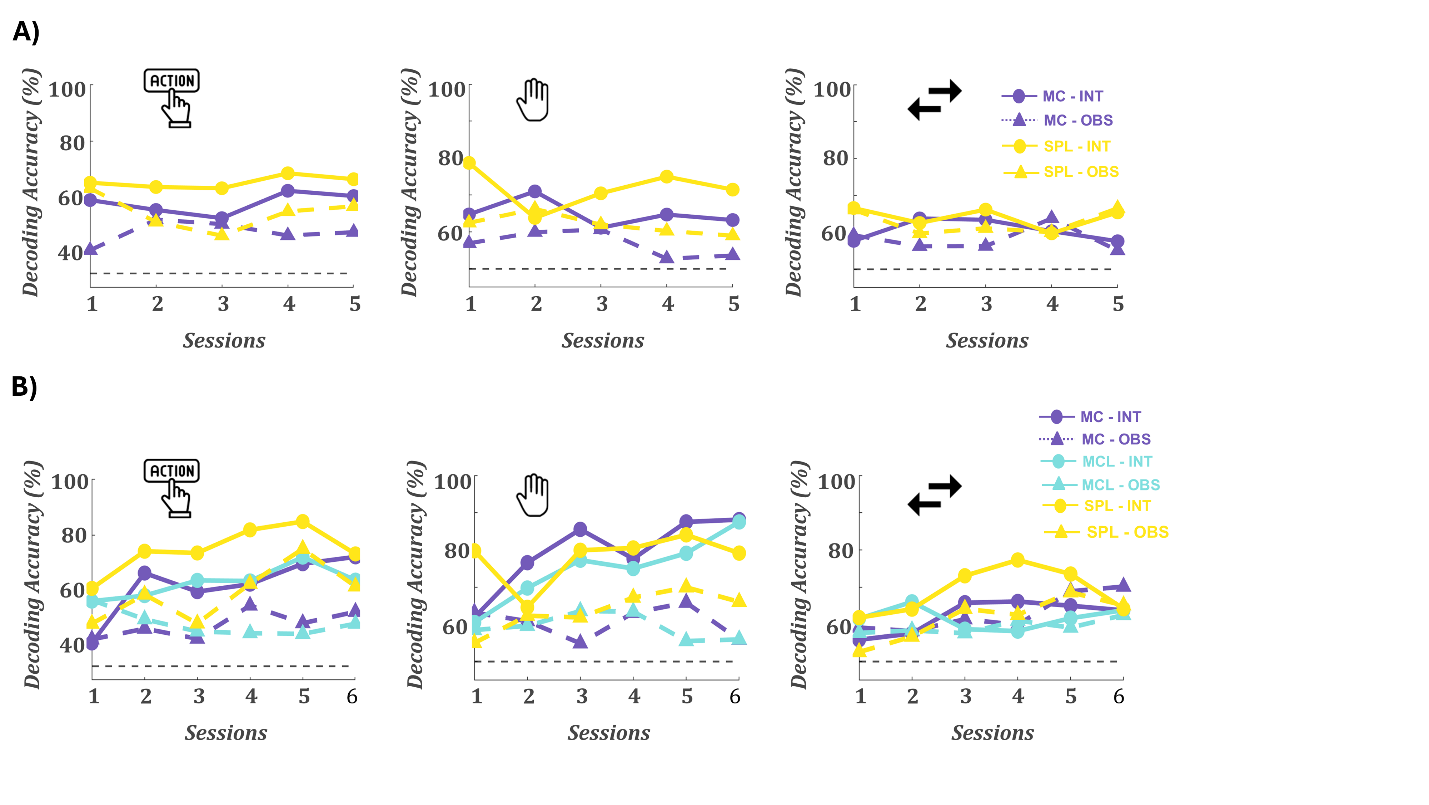

**(A)** Participant JJ. **(B)** Participant RD. Line plots show decoding accuracy for each session across all task variables: action (left), hand (middle), and direction (right). Solid lines and circle markers represent intention; dashed lines and triangles represent observation. Colors indicate recording arrays (see legend). MC/MCM/MCL are shown in purple and cyan tones; SPL is yellow. Action and hand decoding remain consistently above chance across sessions for intention in all regions, indicating stable encoding. SPL shows the highest decoding accuracy for action during observation across nearly all sessions. Direction decoding is consistently poor across all arrays and formats. Overall, decoding performance is stable across sessions, particularly for action and hand representations in intention.

**Figure S7:** **Session-wise decoding accuracy within format.**

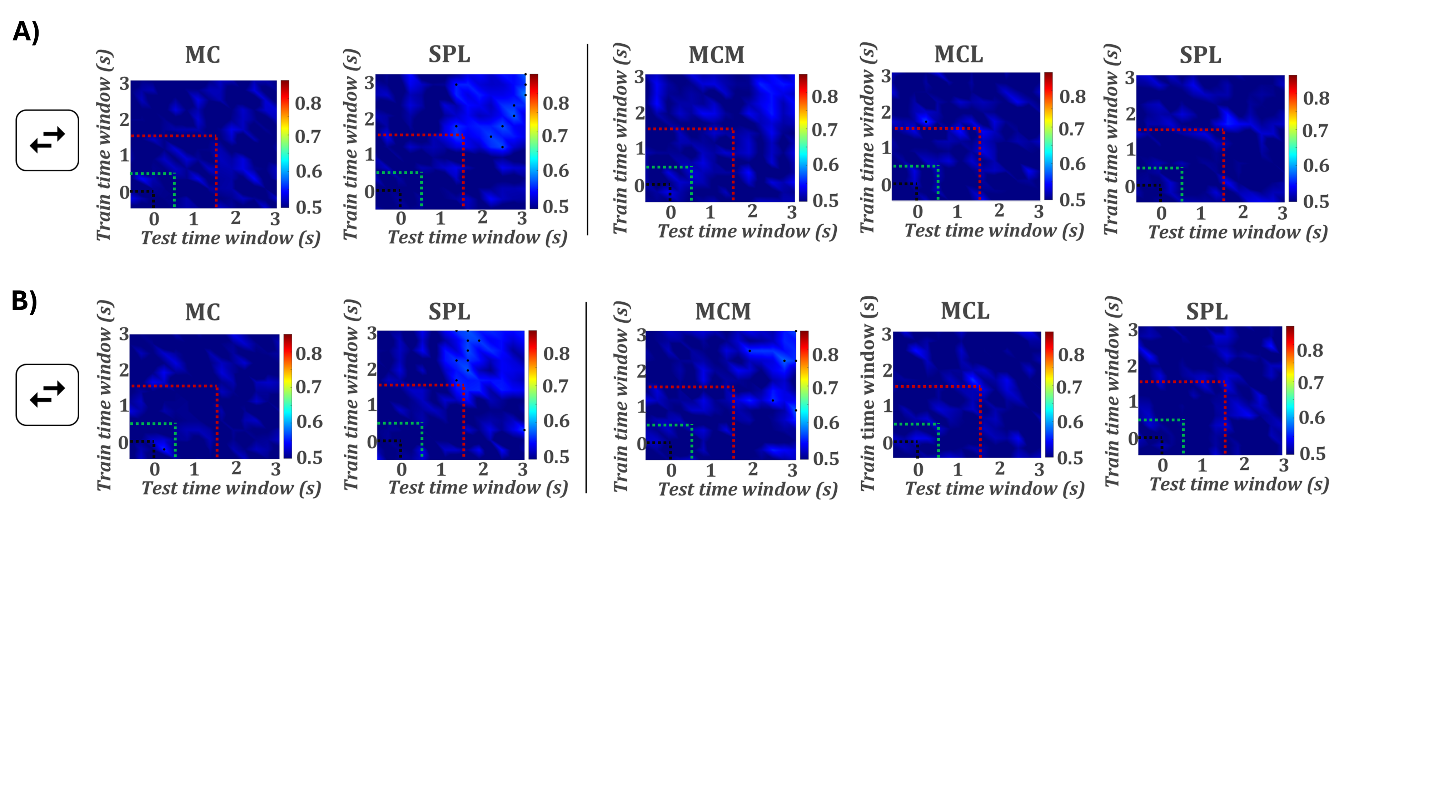

**(A)** Decoders trained on intention and tested on observation. (**B)** Decoders trained on observation and tested on intention. Each heatmap shows decoding accuracy across all combinations of training (y-axis) and testing (x-axis) time bins. Green dashed lines mark the symbolic action cue onset, and red lines the go cue. Dots mark time bin pairs with statistically significant decoding (permutation test). Across all arrays and both cross-decoding directions, decoding performance is low and shows lack of generalization for this task variable.

**Figure S8:** **Additional PCA trajectory comparisons across task variables and control analysis with all units.**

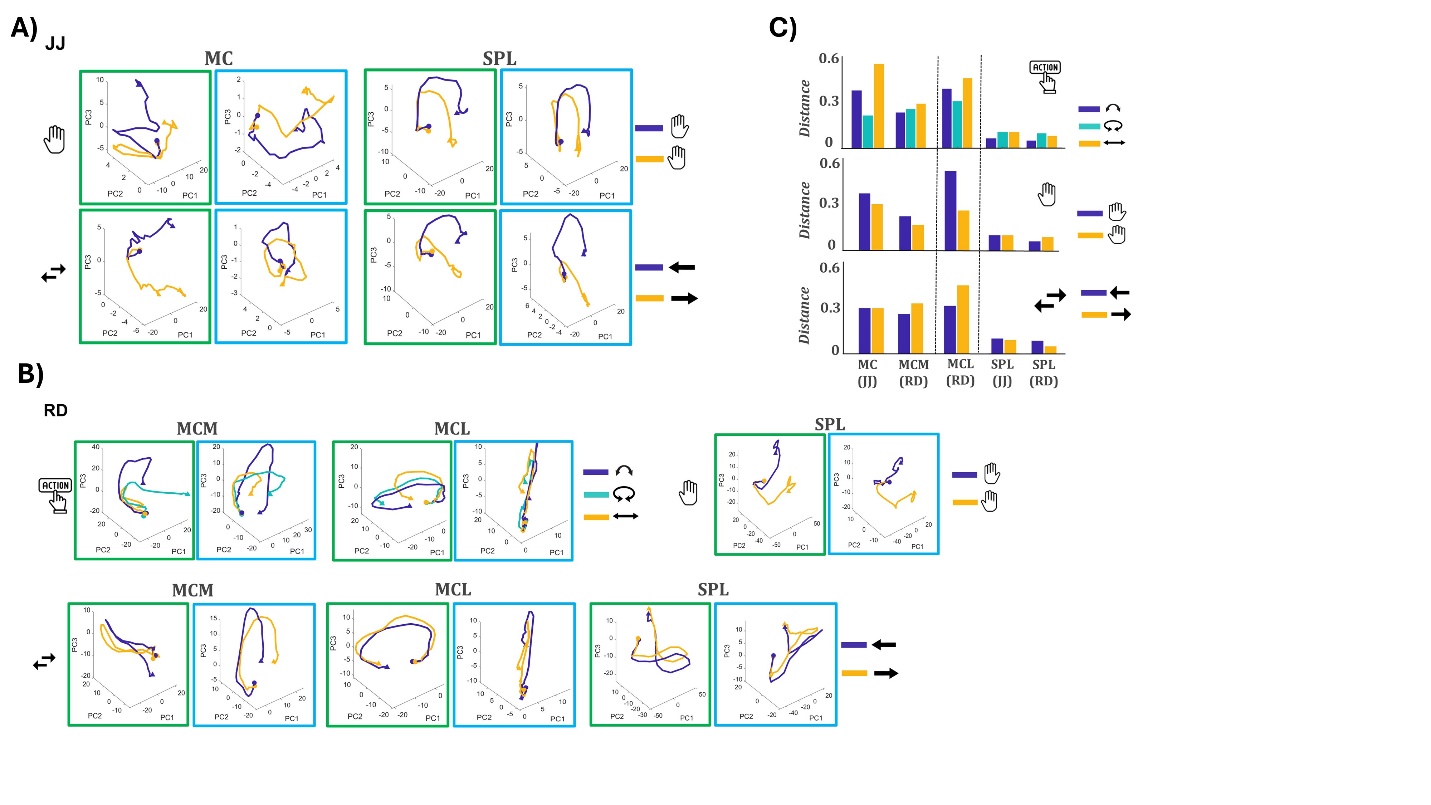

**(A)** PCA trajectories for hand (top) and direction (bottom) in JJ, plotted for MC and SPL. Each 3D plot shows the first three principal components. Within each region, the left panel (green outline) shows intention and the right panel (blue outline) shows observation. Trajectories in MC differ substantially between formats, while SPL trajectories appear very similar for both task variables. (**B)** Same analysis for RD. Shown are action trajectories for MCM and MCL (top row), hand for SPL (top right), and direction for all three arrays (bottom row). Consistent with JJ, trajectories in MCM and MCL differ across formats, while SPL exhibits similar trajectories between intention and observation for both hand and direction. **(C)** Alignment analysis for action, hand, and direction when including all recorded units rather than just task-relevant ones. Results are consistent with the main analysis: SPL shows consistently low alignment distances across all variables (d < 0.2), confirming that the high similarity between formats in SPL is not dependent on unit selection.

**Figure S9: UMAP embeddings for all task variables and cortical arrays.**

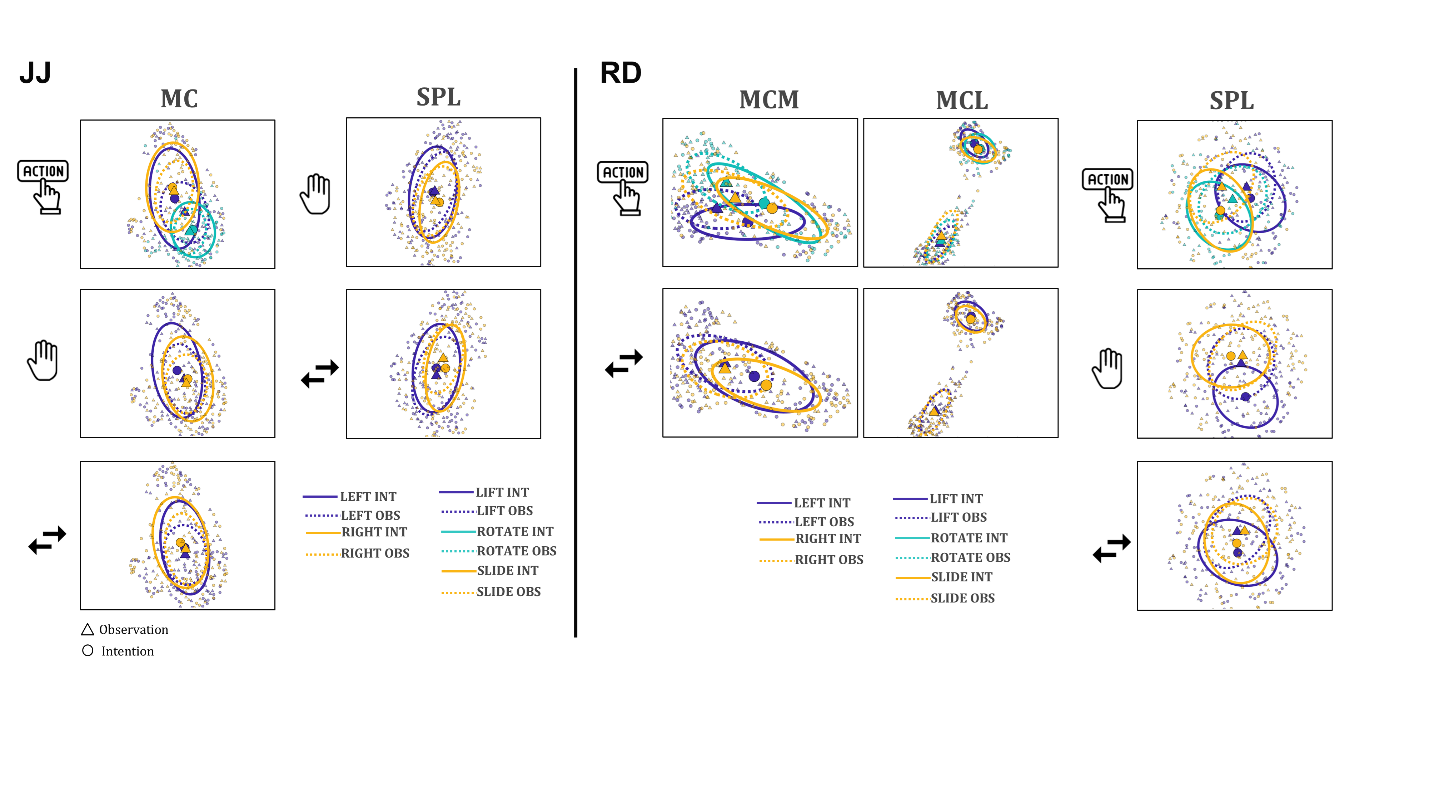

UMAP projections of neural population activity for participants JJ (left) and RD (right), shown separately for action (top row), hand (middle row), and direction (bottom row). Each point represents a trial, with triangles for observation and circles for intention. Ellipses represent 2D distributions per condition (solid for intention, dashed for observation), and colors indicate specific task conditions (e.g., lift, rotate, slide; left/right hand; direction). In MC/JJ, rotation actions show the greatest overlap between intention and observation formats. In MCL/RD, condition clusters are clearly segregated between formats across all variables. SPL exhibits relatively consistent overlap: for the majority of conditions, the centroids of intention and observation responses are close together, suggesting shared structure across formats.

**Figure S10: Within-format decoding of all task conditions using high gamma power**

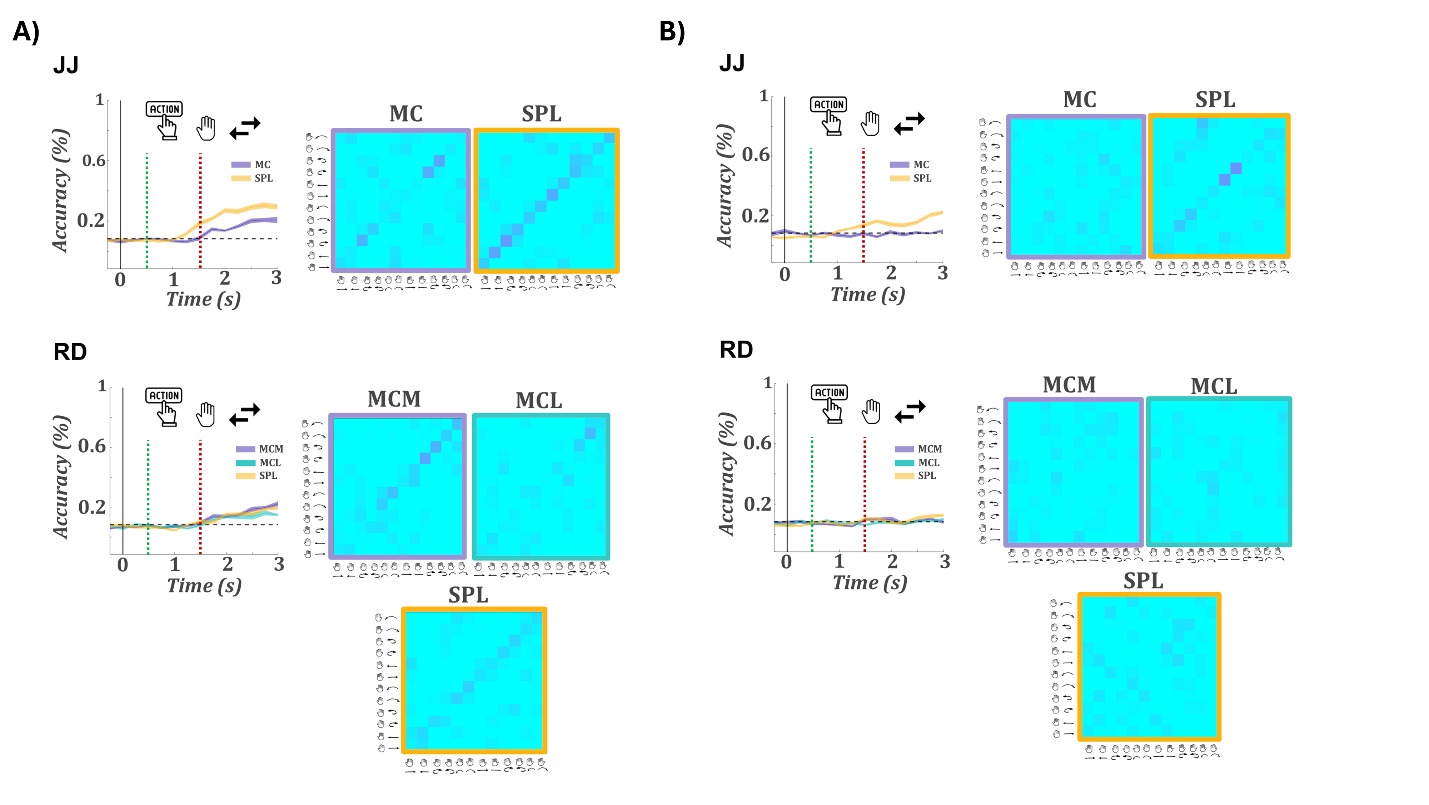

Each panel shows time-resolved decoding accuracy (left) and corresponding confusion matrices (right) for participants JJ (top) and RD (bottom), using high gamma power. Curves represent average decoding accuracy across cross-validation folds; shaded areas indicate ±SEM. Vertical green and red dashed lines mark the symbolic action cue and the go cue, respectively. The horizontal dashed line indicates chance level (1/12 ≈ 8.3%). **(A)** Intention. Decoding accuracy increases after cue onset across all arrays but remains relatively low overall. Confusion matrices show modest condition separability. **B)** Observation. Decoding accuracy remains close to chance in all regions. Only SPL in N1 reaches slightly above-chance performance.

**Figure S11: Cross-format decoding of high gamma activity for all task variables**

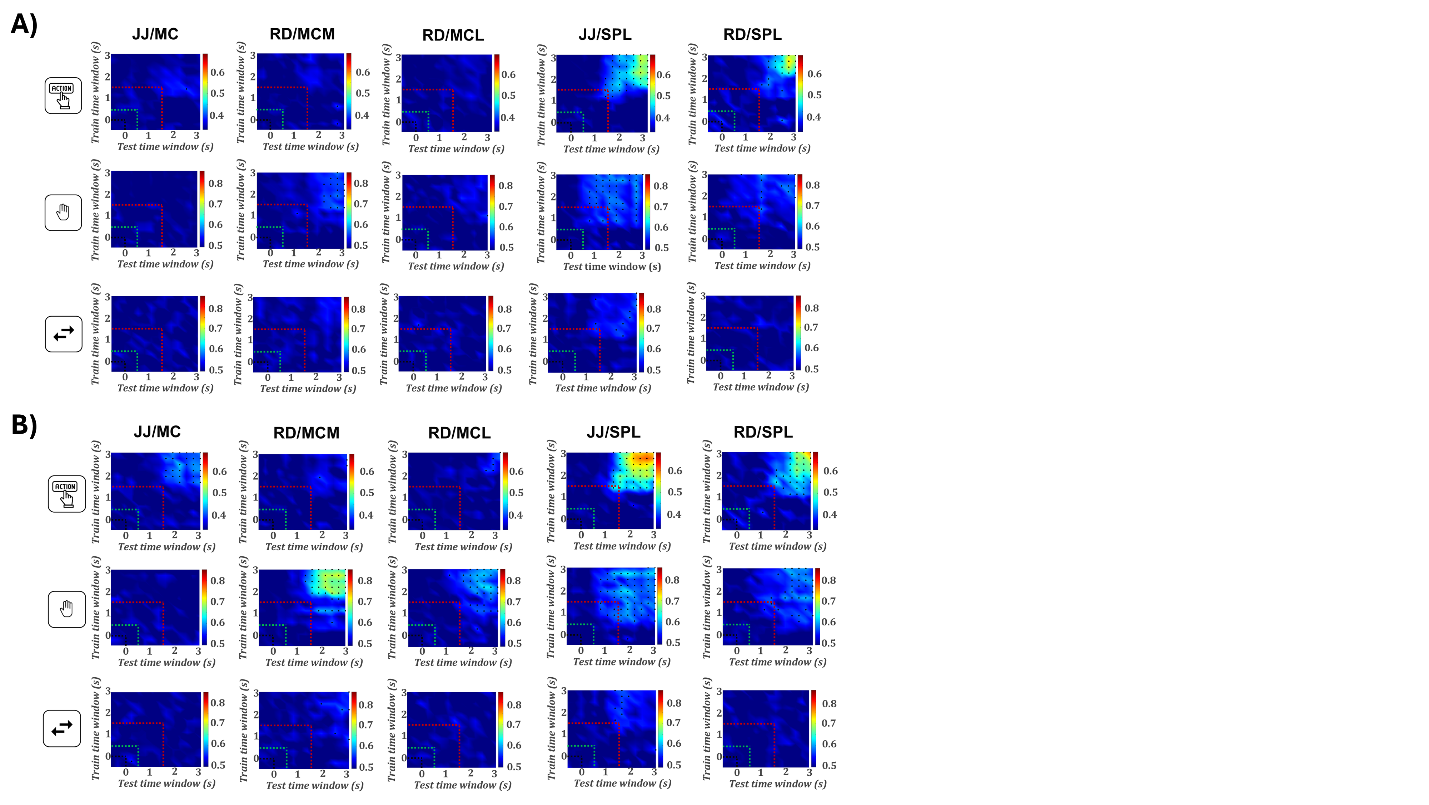

Each panel shows time generalization matrices of decoding accuracy across all combinations of training (y-axis) and testing (x-axis) time bins. Green and red dashed lines indicate the onset of the symbolic cue and the go cue, respectively. Dots mark statistically significant decoding time bin pairs (permutation test). Rows correspond to task variables (top: action, middle: hand, bottom: direction), and columns to arrays from participants JJ and RD. **(A)** Cross-decoding from intention to observation. Only SPL shows clear cross-format decoding for action type in both participants. **(B)** Cross-decoding from observation to intention. SPL again supports action decoding across formats. Additionally, hand can now be decoded in MCM, and weak decoding of action appears in JJ/MC. MCL also shows weak decoding for the right hand.

**Figure S12: Control analyses for congruency in the dissociation tasks.**

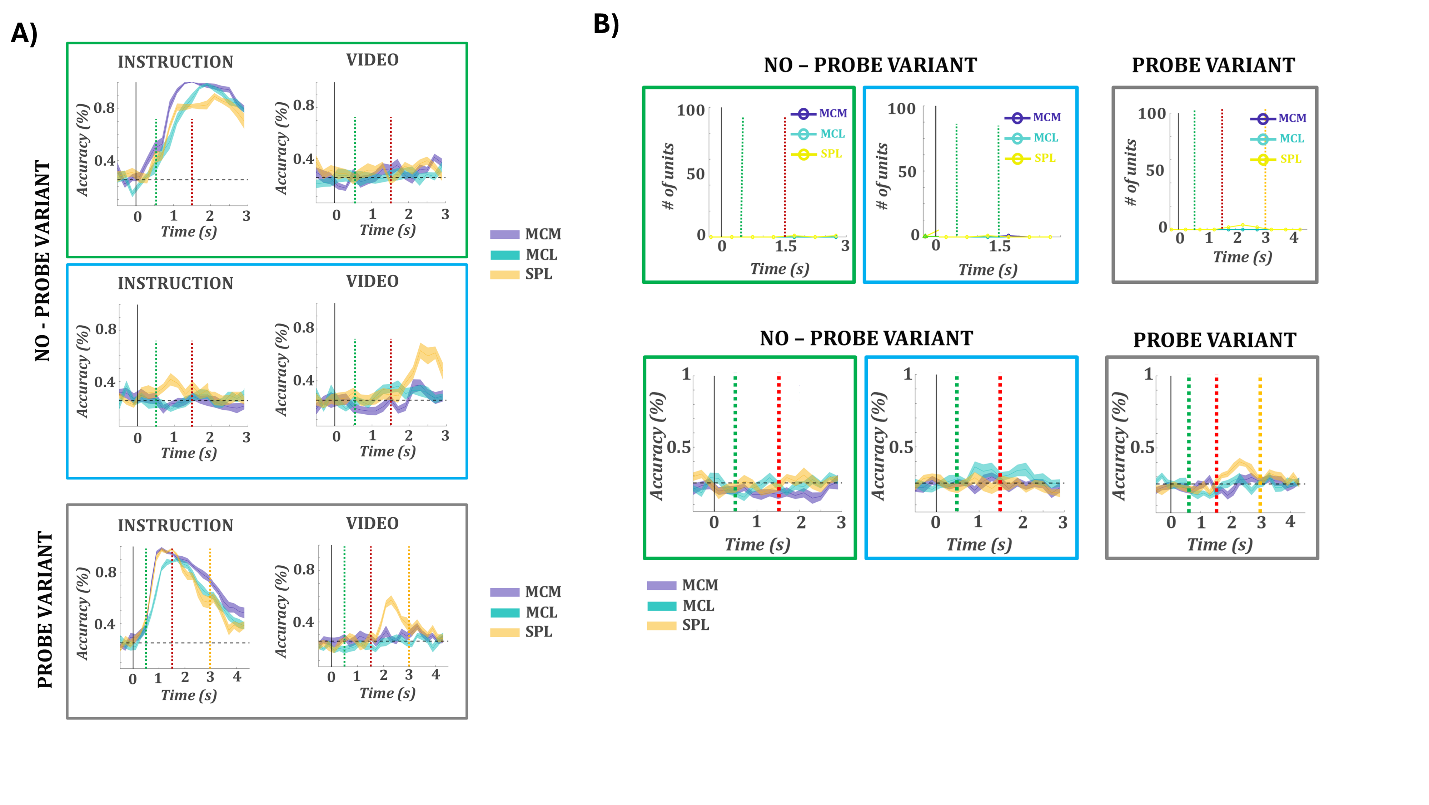

**(A)** Decoding analysis including only incongruent trials from both dissociation tasks. Decoding performance for the instructed (left) and video (right) actions is shown for each variant: no-probe instruction block (green outline), no-probe observation block (blue outline), and probe variant (gray outline). Colored lines indicate mean decoding accuracy over time for MCM (purple), MCL (teal), and SPL (orange); shaded areas show ±SEM across cross-validation folds. The dashed horizontal line marks chance level. Vertical lines denote key task events: hand cue (black), action cue (green), video onset (red), and probe onset (orange). **(B)** Top row: number of tuned units per array for conflict type. Bottom row: decoding accuracy for conflict type. Green outline: no-probe instruction block; blue outline: no-probe observation block; gray outline: probe variant. Shaded areas indicate ±SEM across folds; dashed horizontal line marks chance level; vertical lines as in (A).

**Figure S13: Decoding of all 16 dissociation task conditions.**

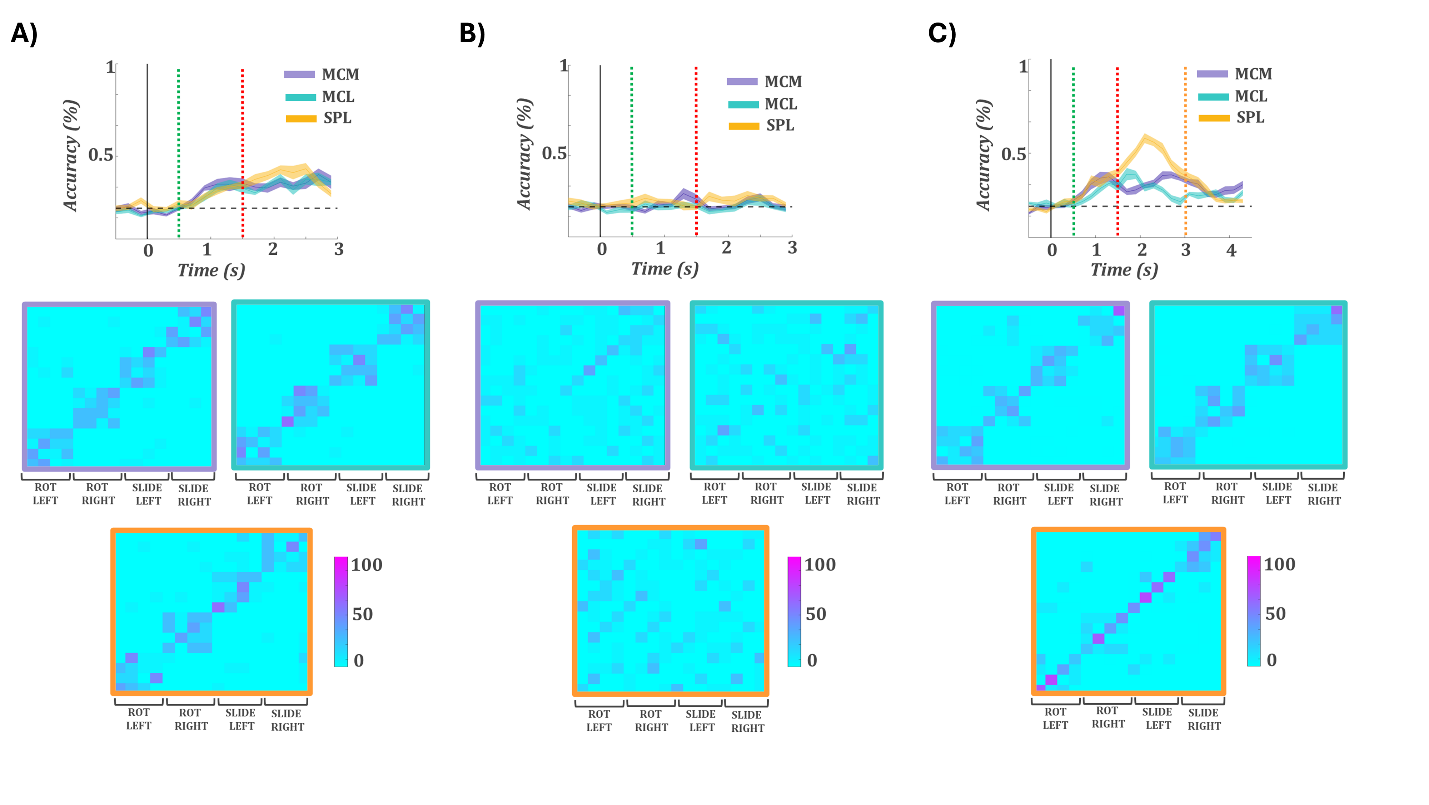

**(A–C)** Time-resolved decoding of the full set of 16 dissociation conditions (2 actions × 2 hands × 4 conflict types) in MCM (purple), MCL (teal), and SPL (orange), shown for the no-probe intention block **(A)**, no-probe observation block **(B)**, and the probe task **(C)**. Line plots show mean classification accuracy over time; shaded areas represent ±SEM across cross-validation folds. The dashed horizontal line indicates chance level (1/16 = 6.25%). Vertical lines mark the hand cue (black), action cue (green), video onset (red), and probe onset (orange). Confusion matrices below each plot show classifier performance at the peak decoding time for each region.

**Tables:**

**Table S1: Experimental summary:** **sessions, trial counts, and recorded units per region**

| **SUBJECT** | **SESSION DATE** | **REPS/CLASS** | **MC** | **SPL** | **MCM** | **MCL** |
| --- | --- | --- | --- | --- | --- | --- |
| **JJ** | 02/06/25 | 12 | 76 | 72 | - | - |
| **JJ** | 02/13/25 | 12 | 75 | 70 | - | - |
| **JJ** | 02/20/25 | 12 | 91 | 62 | - | - |
| **JJ** | 04/03/25 | 12 | 88 | 62 | - | - |
| **JJ** | 04/10/25 | 12 | 91 | 60 | - | - |
| **TOTAL:** |  |  | **421** | **326** |  |  |
| **RD** | 02/06/25 | 12 | - | 110 | 57 | 72 |
| **RD** | 02/13/25 | 12 | - | 79 | 79 | 86 |
| **RD** | 02/20/25 | 12 | - | 82 | 78 | 84 |
| **RD** | 02/27/25 | 12 | - | 86 | 80 | 100 |
| **RD** | 04/03/25 | 12 | - | 92 | 73 | 60 |
| **RD** | 04/10/25 | 12 | - | 83 | 74 | 77 |
| **TOTAL:** |  |  |  | **532** | **441** | **479** |

**Table S2: Explained variance of neural trajectories across regions and formats**

| **ACTION** | **Intention (%)** | **Observation (%)** | **Total (%)** |
| --- | --- | --- | --- |
| **JJ/MC** | 84.07 | 47.7 | 76.2 |
| **JJ/SPL** | 79.2 | 67.1 | 73.8 |
| **RD/MCM** | 68.6 | 59.2 | 66.4 |
| **RD/MCL** | 58.4 | 48.9 | 61.6 |
| **RD/SPL** | 78.2 | 65.2 | 72.8 |
| **HAND** | **Intention (%)** | **Observation (%)** | **Total (%)** |
| **JJ/MC** | 81.7 | 31.3 | 72.6 |
| **JJ/SPL** | 84.6 | 69.5 | 77.6 |
| **RD/MCM** | 93.6 | 64.6 | 88.4 |
| **RD/MCL** | 79.9 | 62.4 | 78.5 |
| **RD/SPL** | 85.7 | 71 | 80 |
| **DIRECTION** | **Intention (%)** | **Observation (%)** | **Total (%)** |
| **JJ/MC** | 79.6 | 34.3 | 71 |
| **JJ/SPL** | 82.5 | 71.8 | 77.1 |
| **RD/MCM** | 71.8 | 61.5 | 71.7 |
| **RD/MCL** | 79.2 | 59.6 | 77.2 |
| **RD/SPL** | 83.4 | 72.8 | 80 |

**Table S3: Dissociation task: sessions, trial counts, and recorded units per region**

| **SUBJECT** | **SESSION DATE** | **REPS/CLASS** | **SPL** | **MCM** | **MCL** |
| --- | --- | --- | --- | --- | --- |
| **RD** | 05/09/25 | 10 | 88 | 55 | 41 |
| **RD** | 05/15/25 | 10 | 94 | 91 | 87 |
| **RD** | 05/23/25 | 10 | 92 | 77 | 70 |
| **RD** | 05/29/25 | 10 | 95 | 81 | 71 |
| **RD** | 06/05/25 | 10 | 80 | 98 | 84 |
| **RD** | 06/06/25 | 10 | 87 | 69 | 55 |
| **TOTAL:** |  |  | **536** | **471** | **408** |

**Table S4: Dissociation task with probe: sessions, trial counts, and recorded units per region**

| **SUBJECT** | **SESSION DATE** | **REPS/CLASS** | **SPL** | **MCM** | **MCL** |
| --- | --- | --- | --- | --- | --- |
| **RD** | 06/12/25 | 15 | 102 | 87 | 90 |
| **RD** | 06/24/25 | 15 | 94 | 83 | 50 |
| **RD** | 07/03/25 | 15 | 116 | 85 | 86 |
| **RD** | 07/23/25 | 15 | 82 | 76 | 72 |
| **RD** | 07/24/25 | 15 | 90 | 83 | 88 |
| **TOTAL:** |  |  | **484** | **414** | **386** |
